# Activation-pathway transitions in human voltage-gated proton channels revealed by a non-canonical fluorescent amino acid

**DOI:** 10.1101/2022.06.30.498345

**Authors:** Esteban Suárez-Delgado, M. E. Orozco-Contreras, Gisela E. Rangel-Yescas, León D. Islas

**Affiliations:** Department of Physiology, School of Medicine, UNAM, Mexico City,04510, Mexico; Department of Biology, Xenon Pharmaceuticals Inc., 3650 Gilmore Way, Burnaby, BC V5G 4W8, Canada

## Abstract

Voltage-dependent gating of the voltage-gated proton channels (H_V_1) remains poorly understood, partly because of the difficulty of obtaining direct measurements of voltage sensor movement in the form of gating currents. To circumvent this problem, we have implemented patch-clamp fluorometry in combination with the incorporation of the fluorescent non-canonical amino acid Anap to monitor channel opening and movement of the S4 segment. Simultaneous recording of currents and fluorescence signals allows for direct correlation of these parameters and investigation of their dependence on voltage and the pH gradient (ΔpH). We present data that indicate that Anap incorporated in the S4 helix is quenched by an aromatic residue located in the S2 helix, and that motion of the S4 relative to this quencher is responsible for fluorescence increases upon depolarization. The kinetics of the fluorescence signal reveals the existence of a very slow transition in the deactivation pathway, which seems to be singularly regulated by ΔpH. Our experiments also suggest that the voltage sensor can move after channel opening and that the absolute value of the pH can influence the channel opening step. These results shed light on the complexities of voltage-dependent opening of human H_V_1 channels.

**Significance statement:** The activation mechanisms of voltage-gated proton channels (H_V_1) are not well understood. Here we have combined patch-clamp fluorometry and a fluorescent non-canonical amino acid to uncover transitions in the activation pathway of human H_V_1 that are modulated by voltage and the pH gradient.

## Introduction

Voltage-gated, proton-permeable ion currents in a large variety of cell types and organisms are produced by the H_V_1 gene (HVCN1 in humans), which encodes a membrane protein that is a member of the superfamily of voltage-sensing domains (VSDs) (Sasaki et al., 2006; Ramsey et al., 2006). These VSDs are also encountered in voltage-sensitive phosphatases (VSP) and voltage-gated ion channels (VGIC), and their principal function is to detect the membrane potential difference and translate it into a conformational change that activates VSP and opens VGIC (Catacuzzeno and Franciolini, 2022).

H_V_1 is thought to form ion channels activated by voltage and employing a mechanism of activation similar to the VSDs of canonical voltage-gated potassium, sodium and calcium channels (Gonzalez et al., 2012). The functions of H_V_1 channels are diverse, including intracellular pH regulation (Ma et al., 2022), charge compensation during immune response (Ramsey et al., 2009), modulation of flagellar beating in spermatozoa (Lishko and Kirichok, 2010), bioluminescence (Eckert and Sibaoka, 1968), and possible roles in calcification processes in marine organisms (Taylor et al., 2011; Rangel-Yescas et al., 2021). Also, H_V_1 is involved in different pathologies such as B cell malignancy (Hondares et al., 2014), breast cancer (Wang et al., 2012), and post-ischemic brain injury (Wu, 2014; Yu et al., 2020); consequently, in recent years H_V_1 has emerged as a possible pharmacological target (Zhao et al., 2018; Zhang et al., 2022).

Among all voltage-gated proton channels sequenced, the human orthologue, hH_V_1, is the most widely studied (Musset et al., 2008). This channel is thought to be a functional dimer formed by two subunits comprising an intracellular N-terminus, a bundle of four transmembrane helices (TMH, S1-S4) in the VSD fold, and a long intracellular alpha helix that mediates a coiled-coil interaction, mainly responsible for dimerization and cooperative activation (Koch et al., 2008; Lee et al., 2008; Tombola et al., 2008). The fourth alpha helix, S4, contains positive charges represented by three arginine residues (R205, R208, and R211) in the characteristic VSD repeats. S4 is thought to undergo an outward displacement and rotation in response to depolarization, mostly in accordance to the helical screw rotation and displacement model of voltage-gating (Li et al., 2015). Unlike canonical VGIC, H_V_1 lacks the two TMH that make up the pore region in canonical ion channels (S5 and S6); therefore, the VSD of H_V_1 has a double function: it is responsible for detecting the electrical potential across the membrane and forming the pathway through which the protons will move once the channel is activated.

A characteristic of H_V_1 activation is its dependence on the pH gradient or ΔpH (pH_o_ - pH_i_). Native proton channels were first shown to open at more negative voltages when the proton gradient points in the outward direction (Cherny et al., 1995). It was shown that for every unit of ΔpH, the voltage of mid activation shifts ~40 mV. Subsequently, all native and cloned proton channels have been found to approximately follow this rule (Sasaki et al., 2006; Ramsey et al., 2006; Rangel-Yescas et al., 2021; Musset et al., 2008; Smith et al., 2011; Zhao and Tombola, 2021). Recent experiments suggest that the proton gradient produces this effect by acting on the voltage sensor and not only affecting a close-to-open transition, since gating currents and channel opening are similarly modulated (Schladt and Berger, 2020; Carmona et al., 2021). However, the molecular mechanisms through which protons modulate voltage-sensor function are not known. Due to technical difficulties, such as not having a pore structure separate from VSD or the impossibility of patch-clamping without protons in the experiments, gating current recordings of H_V_1 channels have been obtained from mutants of the *Ciona* H_V_1 (ciH_V_1) orthologue (Carmona et al., 2018, 2021) or mutants of human H_V_1 (De La Rosa and Ramsey, 2018).

Patch-clamp Fluorometry (PCF) has been used to overcome these difficulties as a powerful tool that allows investigation of electrically silent conformational changes in VSDs associated with channel gating (Kusch and Zifarelli, 2014a), through following fluorescence intensity changes of a dye that generally is attached to the channel protein via chemical modification of cysteine residues or a fluorescent protein genetically encoded (Kusch and Zifarelli, 2014b). In ciH_V_1 and hH_V_1 proton channels, this technique has been employed to obtain evidence of cooperative gating, S1 movement during activation, and the pH sensitivity of S4 movement (Mony et al., 2015; Schladt and Berger, 2020; Qiu et al., 2013). As in other voltage-gated channels, these studies made use of the fluorophore tetramethylrhodamine (TMR), which was tethered to the S3-S4 linker, an extracellular-facing part of the channel (Cowgill and Chanda, 2019). While fluorophores like TMR have provided a means to observe conformational changes in many voltage-gated ion channels, including H_V_1, due to their large size and cysteine reactive nature, they are difficult to incorporate in membrane-embedded portions of channels, without possible large perturbation of the protein fold or result in unspecific incorporation. For these reasons, incorporating a small fluorophore in the middle of transmembrane helices would be advantageous. Here, we exploit the small size, comparable to aromatic amino acids, of a genetically encoded fluorescent non-canonical amino acid, Anap (Chatterjee et al., 2013). By incorporating Anap at various positions in the S4 segment, we assess the conformational changes of the human voltage-gated proton channel (hH_V_1) in response to voltage activation and pH modulation by performing patch-clamp fluorometry (PCF). We find our measurements can resolve a transition during the deactivation process that is strongly modulated by pH. Furthermore, we find that the changes in Anap fluorescence could be partially due to interaction with aromatic amino acids within H_V_1.

## Results

### Incorporation of Anap into hH_V_1

To study voltage-dependent transitions in a voltage sensor using patch-clamp fluorometry, it is desirable that the introduced fluorescent probe does not produce a major structural perturbation of the target protein. The relatively recently developed probe Anap (3-(6-acetylnaphtalen-2-ylamino)-2-aminopropanoic acid) is a small non-canonical fluorescent amino acid (Figure 1B), which has been shown to be easily genetically-encoded in proteins expressed in eukaryotic cells (Chatterjee et al., 2013; Puljung, 2021) Moreover, Anap has been successfully used as a reporter of voltage-dependent conformational changes (Kalstrup and Blunck, 2013) and as a FRET pair (Gordon et al., 2018) to probe ion channel dynamics. In this study, Anap was inserted into specific positions of the S4 segment of the human H_V_1 proton channel (hH_V_1) sequence (Figure 1A), with the purpose of examining its voltage and pH-dependent dynamics.

**Figure 1.**
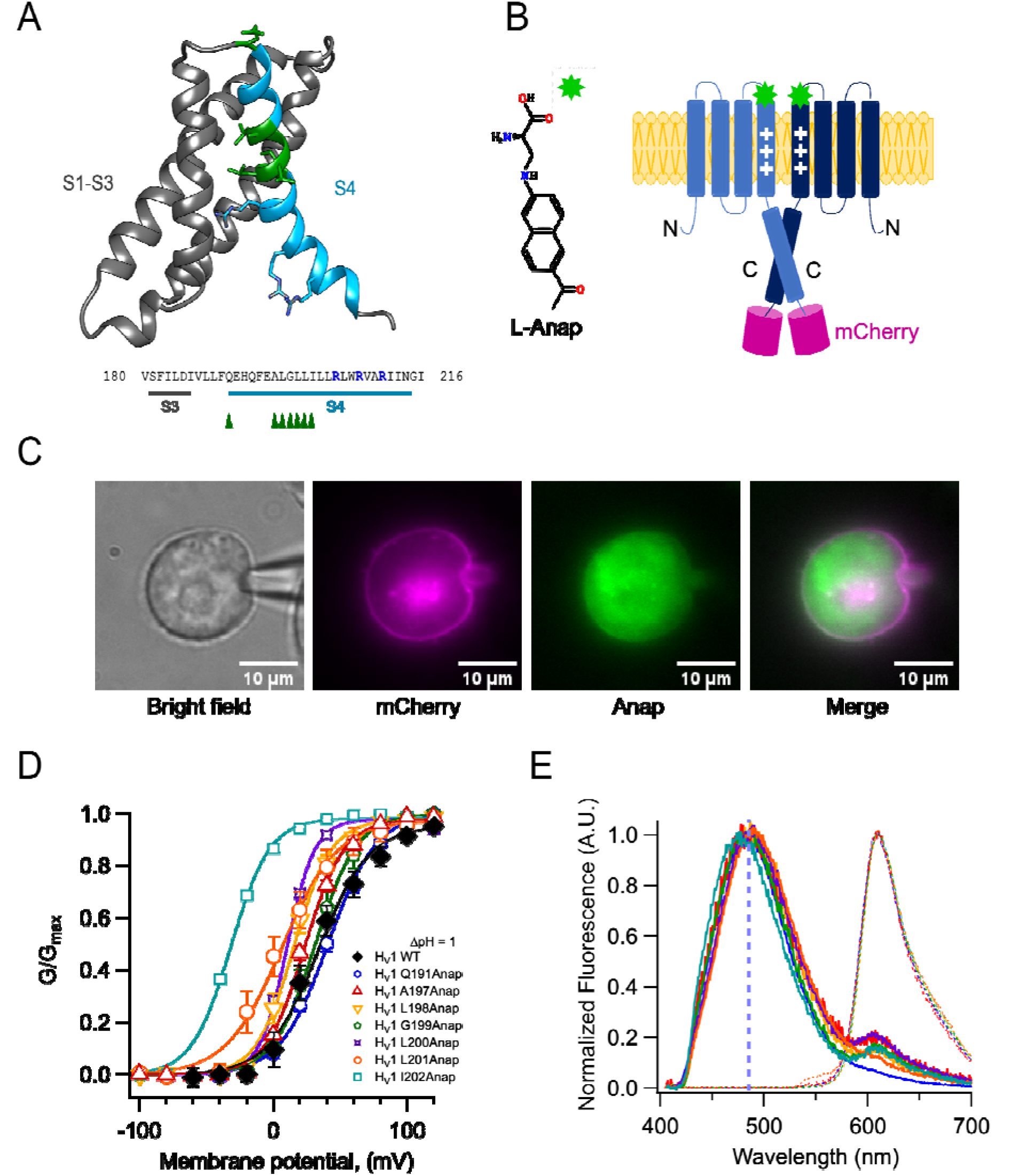
Anap as a fluorescent probe in hH_V_1. A) Ribbon representation of transmembrane segments S1-S4 of closed hH_V_1 based on the model of Randolph et al. (Randolph et al., 2016). S1-S3 are in grey whereas S4 is in light blue. S4 positively charged arginine residues are shown as cyan sticks, whereas the residues where Anap was incorporated individually in the S4 segment are depicted as green sticks and with green arrow heads in the S3-S4 sequence below; positively charged arginine residues are indicated in marine blue. B) Structure of non-canonical amino acid Anap (left), and a schematic representation (right) that shows the incorporation of Anap (green star) into the hH_V_1 dimer expressed in HEK293 cells. An mCherry fluorescent protein (magenta cylinder) was fused to the C-terminal end of hH_V_1 as an Anap incorporation reporter. C) Images of a representative Patch-clamp Fluorometry (PCF) experiment, showing the voltage-clamped cell and the co-localization of Anap and mCherry fluorescence in the cell membrane for Anap incorporated at position Q191 of hH_V_1. D) G-V curves obtained from currents produced by each hH_V_1 mutant rescued by Anap incorporation. All G-V s were obtained at ΔpH=l and compared with hH_V_1 WT. Continuous lines are the fit of the conductance data to equation 1; fit parameters are summarized in Supplemental table I. The incorporation of Anap at the 1202 site sifts the G-V ~65 mV to more negative potentials. Data shown are mean ± s.e.m. E) Normalized mean emission spectrum of Anap (continuous lines) and mCherry (dashed lines) at each incorporation site (color code from D) recorded at resting potential in non-patched cells. Q191(n=15); A197(n=8); L198(n=7); G199(n=6); L200(n=10); L201(n=30); I202(n=10). The vertical blue line indicates the peak emission of Anap in water (486 nm). A second emission peak can be distinguished in every position inside S4 where Anap was incorporated, except Q191Anap. This peak is located around 610 nm which coincides with the peak emission of mCherry.

We selected the S4 helix as insertion target, since this region of the channel is proposed to undergo a voltage-dependent outward displacement that has been previously studied with different approaches, including voltage-clamp fluorometry (13-15).

We were able to successfully substitute amino acids by Anap at positions Q191 in the S3-S4 linker and A197, L198, G199, L200, L201, and I202 in the S4. Anap efficiently rescued expression of channels containing an amber stop codon (TAG) in the selected position, as judged both by appearance of red fluorescence produced by the mCherry fluorescent protein appended in the C-terminus (Figure 1C and Figure 1-supplement 1A) or the appearance of proton currents recorded from HEK 293 cells in the whole-cell patch-clamp configuration (Figure 1D, Figure 1-Supplement 2 and Supplementary Table I).

The substituted channels gave rise to voltage-activated currents, with similar range of activation to WT as judged by their conductance vs. voltage (G-V) curves (except I202Anap channels). Furthermore, that Anap was able to specifically rescue TAG-containing channels was demonstrated by control experiments in cells cotransfected with mutant channels and the pANAP plasmid in the absence of Anap, which showed absence or very small proton currents as compared to cells cultured in the presence of Anap (Figure 1-supplement 1B).The observed fluorescence emission spectrum of the Anap signal present in the membrane (Figure 1C), which presumably originates mostly from Anap incorporated into channels, shows that there are no major or systematic variations on the peak emission wavelength (Figure 1E). The peak emission wavelength for all positions varies, but is near 485 nm, except the most C-terminal and presumably deepest position, I202Anap, which is 477 nm (Figure 1-Supplement 3A). This result suggests that the local environment of Anap in these positions, except 202Anap, is very similar and consistent with a mostly polar environment since the peak emission of Anap in aqueous solution is ~486 nm.

Figure 1E also shows that the emission spectra of positions other than Q191Anap exhibit a small extra peak near 610 nm, that corresponds to the peak emission of mCherry. Since the excitation peak of mCherry is 587 nm (Shaner et al., 2004), we performed experiments with hH_V_1-mCherry channels to measure direct mCherry excitation by our 405 nm laser. The results indicate that this peak is mostly produced by direct excitation of mCherry at 405 nm (Figure 1-Supplement 3B).

### Insensitivity of Anap to pH

In order to use Anap as a reporter of conformational changes in H_V_1 proton channels, and given that these channels are able to change the pH of the surrounding solution (De-la-Rosa et al., 2016; Zhang et al., 2016), we first wanted to validate if this fluorophore is insensitive to pH changes. The amino acid form of Anap that we use is the methyl-ester, which contains amino and carboxy groups. The fluorescence of methyl-ester Anap could be affected by protonation because it could alter electron distribution, for this reason, we reasoned that the best assay to test the pH dependence of Anap fluorescence is to use already incorporated Anap. For this purpose, we used the mutant Q191Anap, which incorporates Anap in the S3-S4 loop, which faces the extracellular solution, even in the deactivated state of the channel (Figure 1A). Since the emission spectrum of Q191Anap channels was measured from non-voltage clamped transfected HEK cells, fluorescence was obtained only from membrane regions to assure that the signal comes mostly from channels exposed to the extracellular solution changes and not channels in intracellular compartments, which will not be exposed to the pH changes.

To ensure that most of the fluorescence is collected from channels in the membrane, the membrane region was identified by the fluorescence of mCherry-containing channels that clearly delineates it (Figure 2-Supplement 1). Spectra were then collected only from this small region of membrane.

These measurements showed that the fluorescence of channel-incorporated Anap is not significantly changed in intensity or shape of the emission spectrum over the pH range 3 to 9 (Figure 2A and B), indicating that this fluorophore is insensitive to pH and that Anap fluorescence should not be affected by local pH changes, which might be produced as a consequence of proton currents.

**Figure 2.**
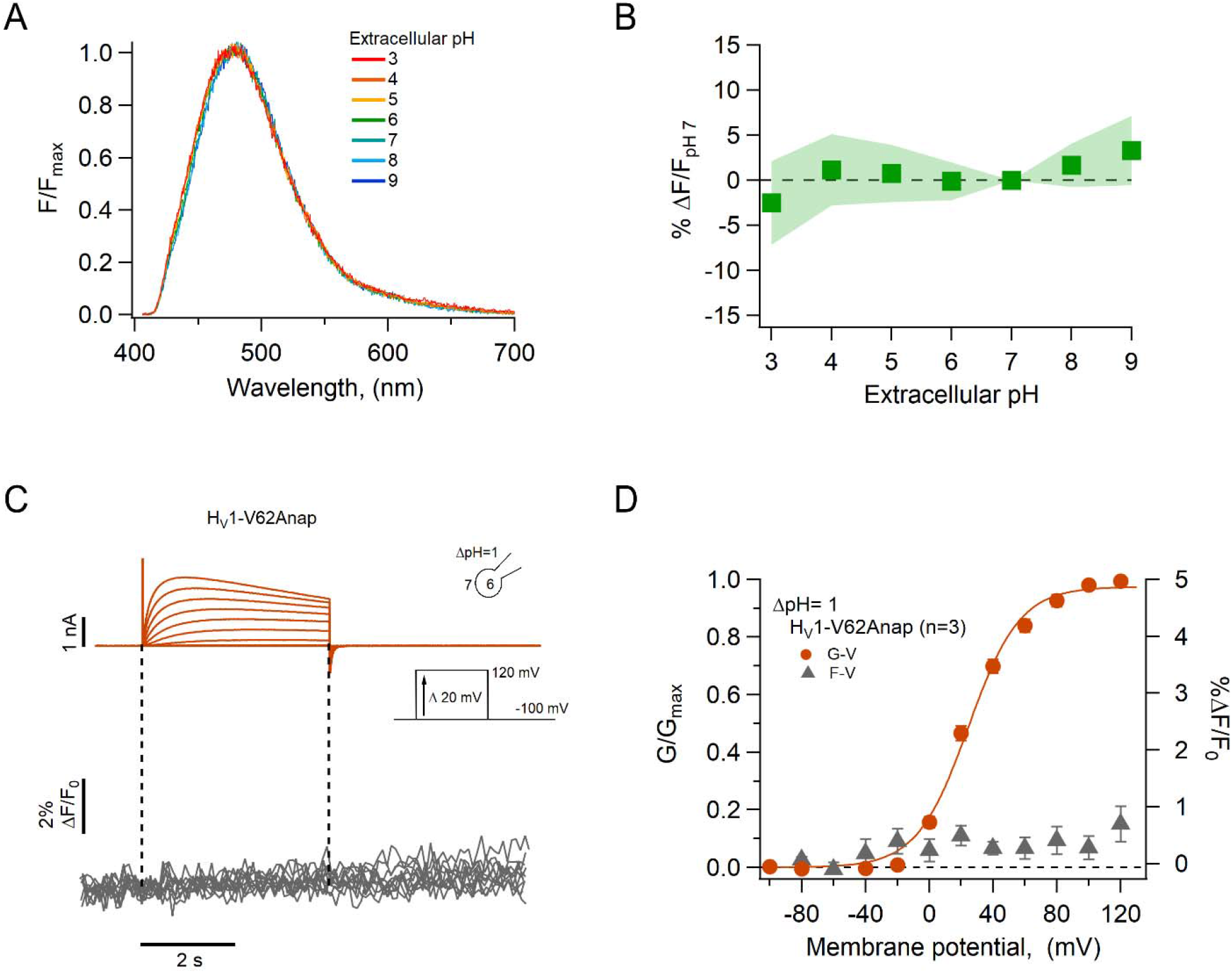
The fluorescence of incorporated Anap is stable to external acidity and local pH changes. A) Mean spectra of Anap fluorescence in the hH_V_1-Q191Anap mutant at each external pH tested (pH_o_) recorded at the resting potential in non-patched cells. The emission peak of spectra of Anap remained inside the wavelength range of 475-480 nm. B) Percentage of fluorescence intensity change normalized to fluorescence at pH_o_ 7 in hH_V_1-Q191Anap mutant (n=13). The intensity was measured from the peak of emission spectra. C) Representative PCF experiments with the hH_V_1-V62Anap mutant. Currents (upper panel, orange traces) and fluorescent signal (lower panel, gray traces) were elicited in response to voltage pulses from −100 mV to 120 mV in steps of 20 mV. D) F-V and G-V relationships from the experiments shown in C. Relative fluorescence changes at the end of voltage test pulses are shown in gray triangles, and conductance is shown in orange circles. The orange continuous line is the fit to equation 1 of G-V data (fit parameters: V_0.5_ = 24.4 ± 1.6 mV; q = 1.5 ± 0.1 e_0_)· Data in B and D are Mean ± s.e.m.

As a further test of our data showing pH-insensitivity of channel-incorporated Anap and to validate the use of Anap in proton channels, we incorporated the amino acid in a position at the N-terminus of the channel, V62Anap. This amino acid is located in the intracellular part of the channel and should be subject to changes in local internal pH during channel activation (De-la-Rosa et al., 2016) but not show changes in fluorescence as a function of voltage-dependent conformational changes. As expected, we did not detect Anap fluorescence changes, although the amino acid was incorporated into functional channels, as judged from proton currents recorded simultaneously with fluorescence (Figure 2C and D). This result further supports the use of Anap in voltage-gated proton channels to measure conformational changes.

### Voltage-dependent changes of Anap fluorescence

Previous experiments in which other dyes like tetra-methyl-rhodamine maleimide (TMRM) were used to label cysteine residues located in the amino-end of the S4 segment of voltage-sensing domains, including the *Ciona* and human H_V_1 channels (Schladt and Berger, 2020; Mony et al., 2015; Qiu et al., 2013), usually result in fluorescence signals that are reduced upon depolarization by a voltage-dependent quenching process (Vaid et al., 2008; Cha and Bezanilla, 1998). In contrast, when we incorporate Anap at position A197, located towards the extracellular end of the S4, depolarization induced an increase of the fluorescence, along with proton currents. The fluorescence increase saturates at depolarized voltages, suggesting that it is produced by a saturable process such as voltage-sensor activation. The direction of this fluorescence change is the same when measured at a ΔpH of 0 or 2, suggesting the same conformational change in the S4 voltage sensor occurs at different pH gradients, albeit over a different range of voltages (Figure 3A and B).

**Figure 3.**
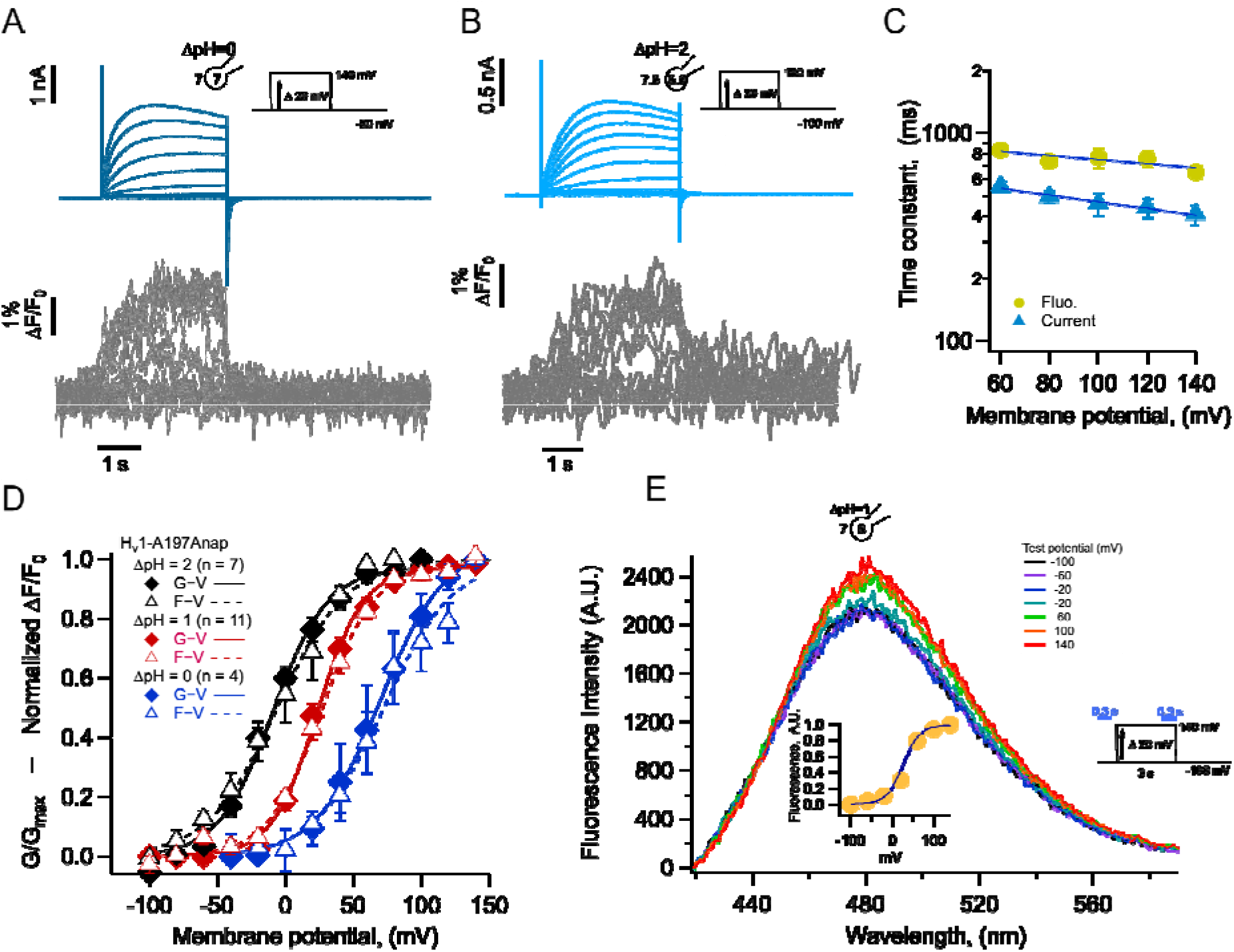
Anap incorporation in position A197 reveals that the movement of S4 is modulated by ΔpH. A-B) Representative PCF experiment with A197Anap at ΔpH=0 and ΔpH=2, respectively. Proton current families (upper panels) are shown in blue traces and fluorescent Anap signal (lower panel) in gray traces. C) Activation time constant of current (blue) and fluorescent (lemon) signals at ΔpH=0 obtained by fitting Eq. 3. The dark blue curve shows the exponential fit to Eq. 4. The fit parameters were: *τ(0)* = 935 ms and *q* = −0.06 e_0_ for fluorescence and 678 ms and −0.09 e_0_ for current. D) F-V (empty triangles) and G-V (filled diamonds) curves and different ΔpH values (ΔpH=0 in blue; ΔpH=1 in red; ΔpH=2 in black). The data were fit to equation 1(G-V, continuous curves; F-V, discontinuous curves) with the following parameters: ΔpH=0; F-V: V_0.5_ = 72.3 ± 6 mV; q = 1.0 ± 0.1 e_0_. G-V: V_0.5_ = 69.6 ± 1.5 mV; q = 1.1 ± 0.1 e_0_. ΔpH =1; F-V: V_0.5_ = 26.6 ± 1.5 mV; q = 1.3± 0.1 e_0_. G-V: V_0.5_ = 23.4 ± 1.3 mV; q = 1.5 ± 0.1 e_0_. ΔpH = 2; F-V: V_0.5_ = −6.1 ± 1.8 mV; q = 1.0 ± 0.1 e_0_. G-V: V0.5 = −9.2 ± 2.3 mV; q = 1.2 ± 2.4 e_0_. E) Emission spectra of Anap in the A197Anap mutant obtained in steady-state (300 ms at the end of holding potential and the end of the test pulse, green bars in the inset) in response to different voltages (color code indicates the test pulse in mV: purple, −60; dark blue, −40; light blue, 20; cyan, 0; light green, 20; dark green, 40; olive, 60; yellow, 80; orange, 100; dark red, 120; red, 140). The inset plots the amplitude of the emission peak as a function of test voltage. The smooth curve is the fit of the fluorescence data at ΔpH=1 shown in panel D. Summary data shown in C and D are mean ± s.e.m.

We compared the kinetics of current and fluorescence by fitting an exponential function to the second half of the signal time course and plotted the time constant as a function of voltage (Figure 3C). Both current and fluorescence have the same voltage dependence, but the current is ~1.3 times faster than the fluorescence. Although not a very large difference, this can be explained by an overestimation of the current time course due to slight proton depletion observed with large currents.

When F-V and G-V curves are plotted together, it is evident that sensor movement paralleled the activation of the proton conductance. At three different values of the pH gradient (ΔpH 0, 1 and 2), both the F-V and G-V curves are almost superimposable and shift along the voltage axis by the same amount of ~40 mV/pH unit (Figure 3D), which is expected of H_V_1 channels (Cherny et al., 1995). Only at ΔpH = 2 the fluorescence signal is shifted to slightly more negative voltages than the conductance and only at voltages at which channel activation begins. The observed voltage shift of the G-V is ~31 mV from ΔpH 2 to 1 and ~43 mV from ΔpH 1 to 0 and is very similar for the F-V curves. This result indicates that the ΔpH-dependence of gating is preserved in channels with incorporated Anap, and that the voltage sensor movement occurs in the same voltage range as the formation of the proton permeation pathway.

Anap is an environmentally sensitive dye, which shifts its emission to red wavelengths in increasingly polar solvents (Figure 3-Supplement 1). To understand the origin of the increased fluorescence observed during activation, we measured the emission spectra of A197Anap in voltage-clamped cells at different voltages. Fluorescence was measured from channels in the membrane region, which are identified by the mCherry signal, similar to Figure 1C. Figure 3E plots the spectra obtained at voltages ranging from −100 to 140 mV and it shows that the fluorescence increases with depolarization and has the same voltage-dependence as the fluorescence measured in Figure 3A, B and C at the same ΔpH (Figure 3E, inset). On the other hand, the peak emission wavelength remains the same at negative or positive voltages, indicating that the increase in fluorescence is not due to depolarization-driven wavelength shifts of the emission spectra. We interpret this result as an indication that Anap incorporated at position A197 remains in a polar environment at all voltages or that small changes in polarity change the quantum yield of Anap but not the emission spectrum.

### H_V_1-197Anap is quenched by a phenylalanine in the S2

The increase of the Anap fluorescence at position 197 in the S4 seen with depolarization could be interpreted as a reflection of an outward movement of the S4 and exposure of Anap to a more polar environment (Figure 3 -Supplement 1), that in principle will produce a red shift of the emission spectrum and an increase of the fluorescence that is detected. However, as shown in Figure 3E, the shape of emission spectrum of Anap remains unchanged at all voltages and with a constant emission peak at ~480 nm, indicating that the fluorophore remains in a polar environment in the closed and open states and thus, a change in local polarity is likely not the principal cause of dequenching.

Many fluorophores can be quenched by aromatic amino acids through mechanisms such as *π*-stacking or photoinduced electron transfer, both mechanisms require close proximity (Islas and Zagotta, 2006; Klymchenko, 2017; Pantazis and Olcese, 2012; Young and Artigas, 2021). Evidence for the existence of a quenching group near Anap incorporated in the S4 can be obtained from examination of the ratio of fluorescence of Anap and mCherry in channels as a function of its position along the S4 (Figure 4-Supplement 1). This analysis shows that for positions deeper into the S4 segment, the Anap/mCherry ratio diminishes, suggesting that in those sites Anap is closer to a quenching group.

For these reasons, it is conceivable that the Anap quencher in H_V_1, could be an aromatic residue located near the introduced fluorophore in the closed state and upon S4 movement, increases its distance, generating the observed dequenching. We used a structural model of H_V_1 derived from experimental data (Randolph et al., 2016) and replaced A197 with Anap. Figure 4A shows Anap in salmon-colored spheres and highlights aromatic residues within the transmembrane domains of a monomer as dotted spheres. A possible candidate for an Anap quencher is F150 (yellow spheres), because this residue is the closest aromatic to Anap that is not in the S4 and F150 will presumably remain in its position as 197Anap undergoes an outward displacement with depolarization. In contrast, other aromatic residues which are closer to 197Anap and are part of the S4, will presumably move with all the S4 as a rigid body. Incidentally, an equivalent phenylalanine to F150 has been identified as the charge transfer center in canonical voltage-gated potassium channels and in H_V_1 (Tao et al., 2010; Hong et al., 2013).

**Figure 4.**
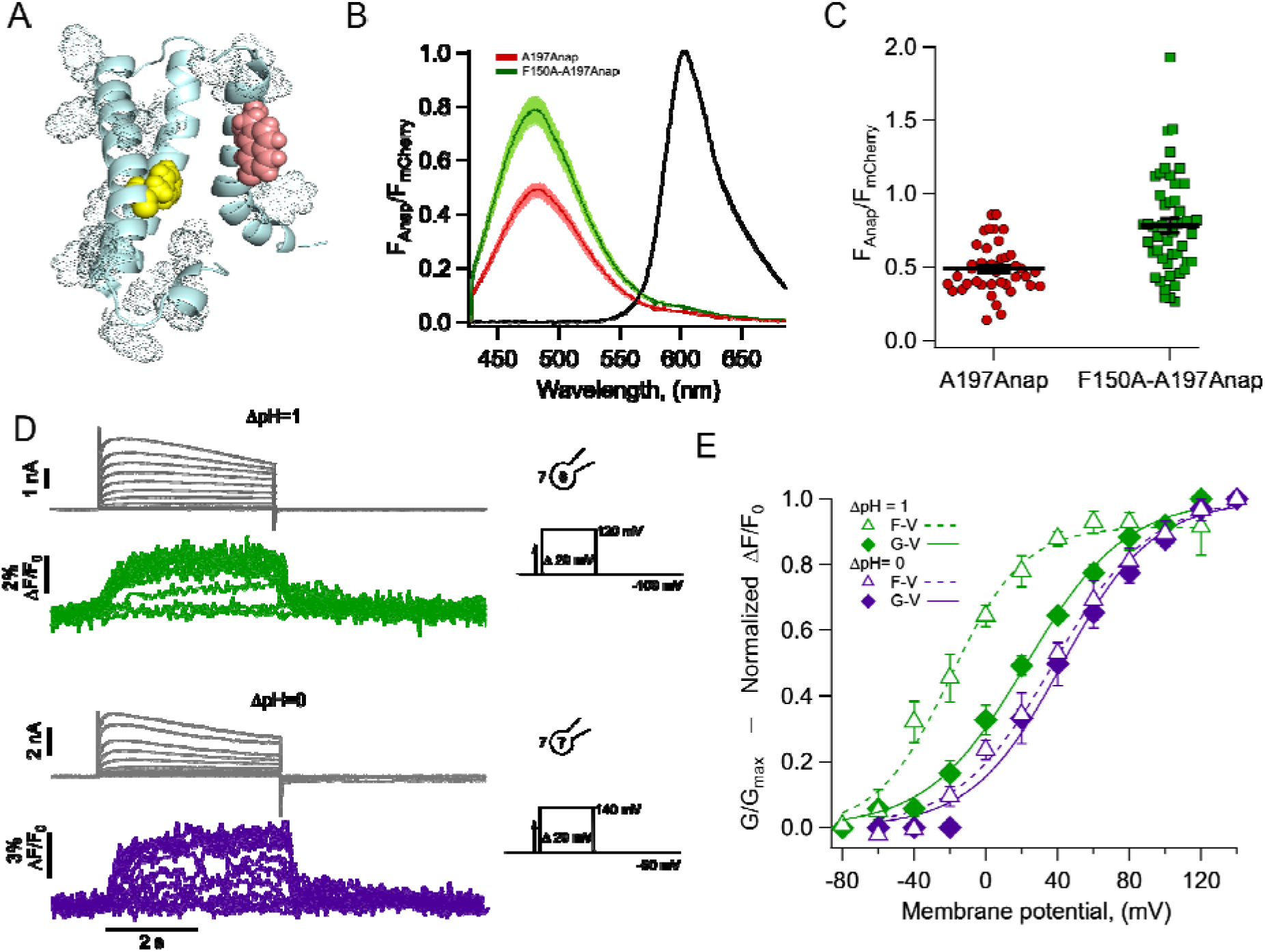
The charge transfer center (F150) is an Anap quencher. A) Cartoon showing the presence of aromatic residues in hH_V_1 (rendered as space-filling dots, main chain in light blue, S3 was removed for illustration). F150 in yellow and Anap in pink. B) Averages of spectra of Anap incorporated in both mutants (H_V_1-A197Anap, red; H_V_1-F150A-A197Anap, green) normalized to the fluorescence of mCherry (black). Spectra were obtained at the resting potential in non-patched cells. The double mutant’s brightness is approximately 60% higher. Shadows represent s.e.m. C) Comparison of the intensity of the emission spectrum peak of Anap normalized to the intensity of the fluorescent protein mCherry between the mutant H_V_1-A197Anap-Cherry (0.49 ± 0.03) and double mutant H_V_1-FI50A-A197Anap-Cherry (0.79 ± 0.05), taken at 48 hours post-transfection. Each point indicates an individual spectrum measured from a single cell; n = 41 and 49, respectively. Black horizontal lines are the mean ± s.e.m. T-test value p <0.001. D) Representative current and fluorescence traces from PCF experiments of the double mutant H_V_1-F150A-A197Anap at ΔpH=1 (upper panel) and ΔpH=0 (lower panel). E) Comparison of G-V (diamonds) and F-V (triangles) relationship between both ΔpH conditions (ΔpH=1 in green; ΔpH=0 in purple) of the double mutant H_V_1-F150A-A197Anap. F-V curve of H_V_1-F150A-A197Anap at ΔpH=0 is shifted negatively around 58 mV compared to ΔpH=1. Boltzmann fit parameters of H_V_1-F150A-A197Anap were: ΔpH=l F-V: V_0.5_=−19.8 ± 2.7 mV; q =1.2 ± 0.1 e_0_; G-V: V_0.5_ = 22.7 ± 2.3 mV; q = 0.9 ± 0.1 e_0_. ΔpH=0 F-V: V_0.5_=38.0 ± 3.0 mV; q =0.9 ± 0.1 e_0_; G-V: V_0.5_ = 42.6 ± 3.8 mV; q = 1.0 ± 0.1 e_0_. Data shown in B, C and E are mean ± s.e.m.

To test this hypothesis, we made the double mutant F150A-A197Anap and estimated the relative amount of basal Anap quenching, by comparing the emission spectra of both Anap and mCherry in the same membrane region. Figures 4B and C shows that the double mutant displays a significantly increased Anap fluorescence (~60 %) relative to mCherry, when compared to A197Anap alone, suggesting that indeed, phenylalanine 150 is capable of quenching Anap in the closed state (at the resting potential of HEK cells of −20 to −40 mV (Thomas and Smart, 2005b) and at the employed ΔpH ~ −0.2 (pH_o_ = 7) most channels should be in the closed state).

Despite having removed the quenching group, F150A-A197Anap channels still show voltage-dependent fluorescence changes (Figure 4D), suggesting the presence of additional quenchers or that in the absence of F150, Anap at 197 becomes sensitive to polarity changes.

The voltage dependence of the fluorescence signals from F150A-A197Anap channels shows significant differences from those of A197Anap alone (Figure 4E). At values of ΔpH of 0 and 1, fluorescence precedes the increase in conductance, indicating that the conformational change of the S4 segment occurs at more negative voltages than the formation of the proton-permeable pathway. This effect is more pronounced at ΔpH = 1. Interestingly, the difference of V_0.5_ of the F-V curve between ΔpH= 0 and 1 is ~58 mV, similar for A197Anap, which is ~46 mV.

### A distinct gating transition detected by Anap fluorescence

The fluorescence time course of the F150A-A197Anap channels shows an interesting characteristic; the OFF signals (F_off_) that are produced at the return of the voltage to the holding potential and represent the return of the voltage sensor to the resting states, show a two-component kinetic behavior. This is particularly evident at ΔpH = 0 (Figure 4D), where F_off_ shows a very rapid quenching followed by a much slower component, suggesting that the voltage sensor can move back to its resting position at varying rates.

To explore the kinetics of fluorescence signals during repolarization, and since this double mutant removes a quenching group, we used the hH_V_1-201Anap channels. We reasoned that this mutant channel, which has Anap in a deeper position in the S4 and presumably closer to F150 in the closed state, might be a better reporter of the kinetics of S4 movement.

Figures 5A, B and C show simultaneous current and fluorescence recordings from hH_V_1-1201Anap channels at three different ΔpH values of 0, 1 and 2. As with the hH_V_1-A197Anap construct, the voltage-dependence of the conductance and fluorescence are almost superimposable and shows a large shift of >40 mV/pH unit (Figure 5D).

**Figure 5.**
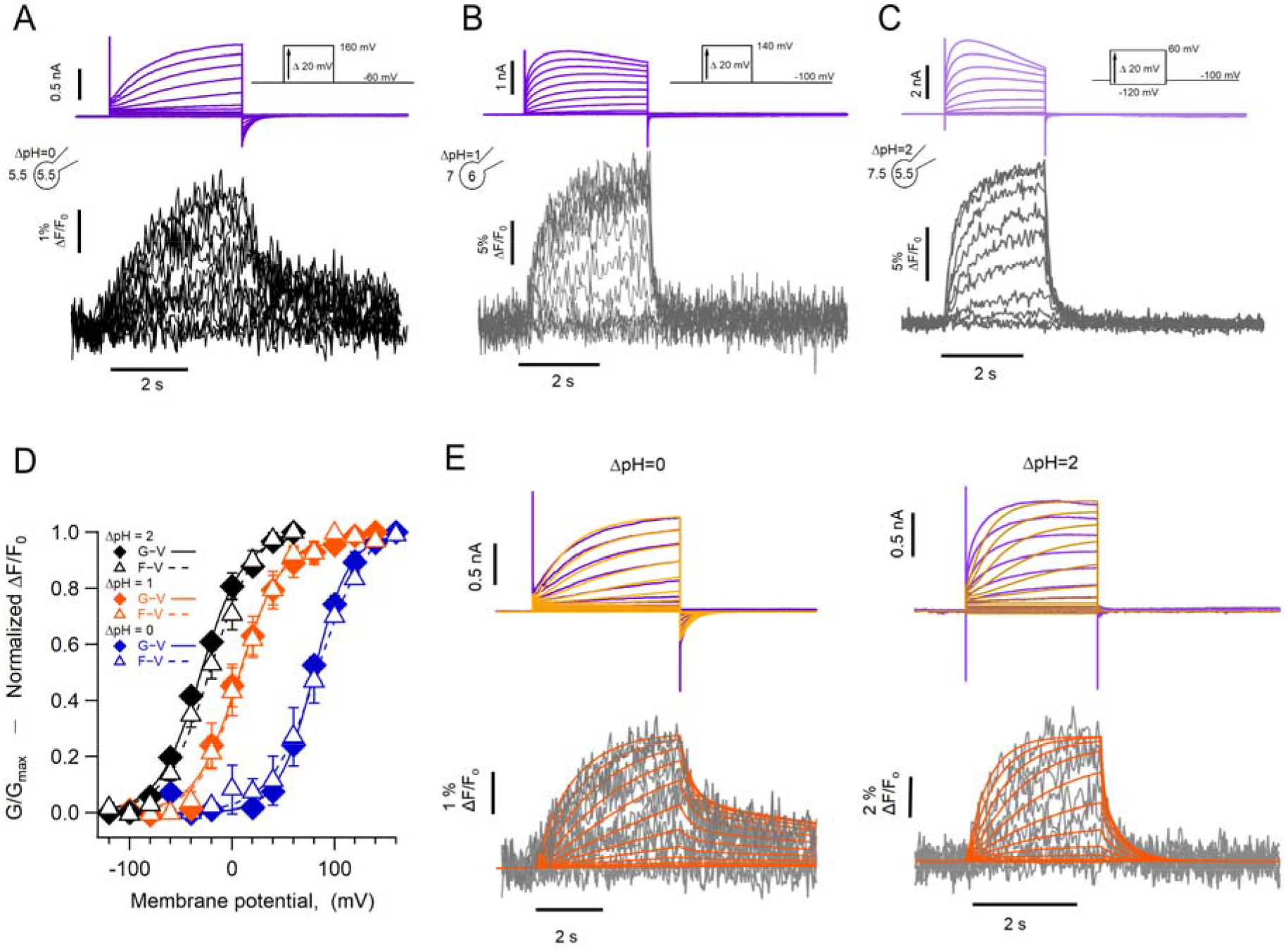
The kinetics of fluorescent signal during deactivation is strongly modulated by pH. Representative PCF experiments with the hH_V_1-L201Anap mutant at: A) ΔpH=0. B) ΔpH=1. C) ΔpH=2. Current families are shown in the upper panel (purple traces) and fluorescent signals in the lower panel (black and gray traces). D) G-V (filled diamonds) and F-V (empty triangles) relationships at ΔpH=0 (blue markers, n = 3), ΔpH=1 (orange markers, n=4) and ΔpH=2 (black markers, n=5) of mutant hH_V_1-1201Anap. Data are mean ± s.e.m. Note that the difference between the activation at ΔpH=1 and ΔpH=0 is around 77 mV/ΔpH unit. Boltzmann fit parameters: ΔpH=0, F-V; V_0.5_= 84.6 ± 2.1 mV, q =1.0 e_0_ ± 0.1. GV; V_0.5_= 79.7 ± 1.8 mV, q = 1.4 ± 0.1 e_0_. ΔpH =1, F-V; V_0.5_= 7.7 ± 1.6 mV, q =1.2 ± 0.1 e_0_. G-V: V_0.5_= 6.3 ± 2.2 mV; q = 1.2 ± 0.1 e_0_. ΔpH = 2, F-V: V_0.5_= −21.1 ± 2.3 mV; q =1.1 ± 0.1 e_0_. G-V: V_0.5_= −30.7 ± 1.9 mV; q = 1.2 ± 0.1 e_0_. E) Comparison of the current and fluorescence at two values of ΔpH with the predictions of the sequential activation model in Scheme I. Experimental current and fluorescence traces are color coded as in A). Simulated current traces are mustard colored and fluorescence traces are orange. Simulation parameters can be found in Supplementary Table II.

The most remarkable feature of these fluorescence traces is that, at ΔpH = 0, the OFF signal during repolarization (F_off_) has two distinct kinetic components. The deactivation tail currents at −60 mV decay exponentially, with a time constant of 141 ±55 ms, while the F_off_ can be fit to a sum of two exponentials with time constants of 129 ± 68 ms and 8.6 ± 0.74 s. (Figure 5-Supplement 1). The presence of the two components in F_off_ suggest that the return of the voltage sensor to its resting state can occur at varying rates. In particular, the slow component is consistent with the immobilization of the off-gating charge observed in monomeric *Ciona* H_V_1 channels (Carmona et al., 2018). The slow off-component is also present at ΔpH = 1 and 2, although its amplitude is smaller. We did not undertake a systematic kinetic analysis of current and fluorescence traces during channel activation due to the alterations that proton depletion cause on the current time course, especially for the larger currents observed at ΔpH = 2.

Instead, to qualitatively understand the kinetics of the fluorescence signals, we used a simplified kinetic model of channel activation (Scheme I), similar to a model that was previously used to study the voltage-dependent kinetics of hH_V_1 (Villalba-Galea, 2014).

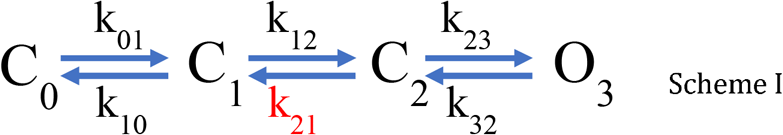

In this model, one of the backward transitions (k_21_) between closed states is set to be much slower than the open to closed transition (O_3_->C_2_), resulting in a large difference between the closing kinetics of the ionic current, mostly determined by k_32_, and the fluorescence signals. This kinetic difference can account for the biphasic behavior of the F_off_ signal, and especially the slow component of its time course (Figure 5E). The model also indicates that when the internal pH is lower than the external pH (ΔpH=2), this slow rate constant is more affected than any other, indicating a conformational step that is especially sensitive to pH.

While the simple model in Scheme I can account qualitatively for the observed kinetics of 201Anap channels, the experimental F-V relationship is superimposable on the G-V curve (ΔpH=0 and 1) or positively shifted by ~10 mV (ΔpH=2) with respect to the G-V curve, which is not a feature predicted by Scheme I and is reminiscent of channels that can open to multiple open states, without the need of full voltage sensor activation (Stefani et al., 1997). This observation suggests that hH_V_1 channels operate via a more complicated mechanism that the sequential gating illustrated by Scheme I, which might include channel opening before complete voltage-sensor movement. We tested a simple version of such an allosteric model and show that it can account, at least qualitatively, for current and fluorescence kinetics and for the relationships between G-V and F-V curves at varying ΔpH (Figure 5-Supplement 2). Interestingly, in this model the slow deactivation rate constant is also the step with the most sensitivity to pH (Supplementary Table III).

### Absolute pH values are determinants of voltage sensor movement

One of the most intriguing characteristics of H_V_1 channel gating, is its steep modulation by the pH gradient. While it has been shown that this modulation depends on the value of ΔpH, regardless of how it is set up (Cherny et al., 1995), there is evidence that the absolute value of pH can also exert an effect on gating (Cherny et al., 2015). In most of our experiments, the pH gradient was set up with a low value of intracellular pH, between 5.5 and 6.0. To test the effect of absolute pH, we carried out experiments with the same ΔpH of 0, with symmetric low (5.5_/_5.5) or high (7_/_7) intra/extracellular pH. The expectation was that, if pH gating of hH_V_1 depends only on the pH gradient, the voltage sensor should move with essentially the same characteristics. Surprisingly, the fluorescence signals display important differences, as do the proton currents. Our results in Figure 6 show that when compared to ΔpH = 0 (5.5_/_5.5), the fluorescence in symmetric pH_o_ and pH_i_ = 7.0 has a rapid return of the F_off_ signal (Figure 6A and B). Interestingly, the voltage dependence of the F-V relationship is very similar for (5.5_/_5.5) or (7_/_7) conditions, while in (7_/_7) the proton current appears at more negative voltages than the bulk of the fluorescence (Figure 6C). These results suggest that the voltage range of movement of the voltage sensor, as reported by the fluorescence of 201Anap, is dependent on the ΔpH, since the V_0.5_ of the F-V is the same in pH_o_/pH_i_ =5.5_/_5.5, while the opening of the proton conduction pathway in pH_o_/pH_i_ =7_/_7, can occur after only a fraction of the voltage sensor movement has occurred and this coupling between voltage sensing and channel opening can be increased by low pH.

**Figure 6.**
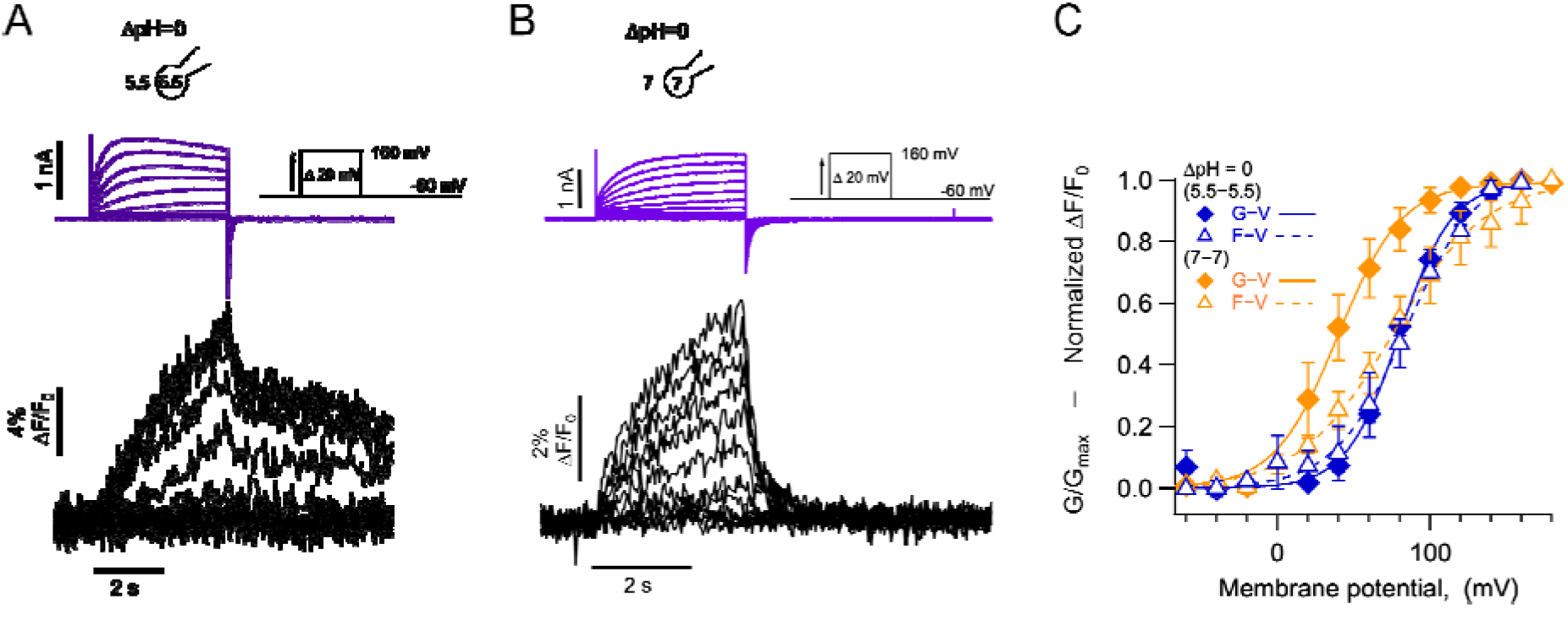
Absolute pH values are gating determinants in hH_V_1-1201Anap. A) Representative PCF experiment at ΔpH=0 (5.5_o_-5.5_i_). Currents are purple and fluorescence black. B) Similar experiment to A) with ΔpH=0 (7_o_-7_i_). Current and fluorescence traces color coded as in A). C) G-V (filled diamonds) and F-V (empty triangles) curves at ΔpH=0 but with different absolute pH values (pH_o_/pH_i_ =5.5_/_5.5 in blue; pH_o_/pH_i_ =7_/_7 in orange, n= 4). Boltzmann fit parameters were pH_o_/pH_i_ =7_/_7 F-V: V_0.5_= 75.3 ± 2.2 mV; q =0.8 ± 0.04 e_0_. G-V: V_0.5_= 39.6 ± 1.3 mV; q = 1.2 ± 0.1 e_0_. pH_o_/pH_i_ =5.5_/_5.5 F-V: V_0.5_= 84.6 ± 2.1 mV; q =1.0 e_0_ ± 0.1. G-V: V_0.5_= 79.7 ± 1.8 mV; q = 1.4 ± 0.1 e_0_. Data are mean ± s.e.m.

## Discussion

In the experiments described here we have implemented patch-clamp fluorometry in combination with incorporation of a fluorescent non-canonical amino acid (NCAA) to study voltage-dependent gating in hH_V_1 proton channels. Although voltage-clamp fluorometry (VCF) using TMRM has been used previously to study H_V_1 channels (Qiu et al., 2013; Mony et al., 2015; Schladt and Berger, 2020), employing the fluorescent Anap NCAA has the advantages of being a smaller size probe and improving the specificity of fluorescence signals, since it is genetically encoded. The small size of Anap allowed us to incorporate the fluorophore into functional channels in several sites along the S4 and the S3-S4 loop. Since Anap was developed as an environmental sensitive probe, the fact that the emission spectrum of Anap in these sites is very similar to that of Anap in water, suggests that these residues are solvated in the native H_V_1. The only position that shows a blue-shifted Anap spectrum is I202, which is the most C-terminal residue explored and might burry the Anap R-group in a more hydrophobic environment.

Since the activity of H_V_1 proton channels can change the local concentration of protons near the conduction site and fluorescence probes have been used to detect these proton fluxes (De-la-Rosa et al., 2016; Zhang et al., 2016), we addressed whether Anap could change its fluorescence as a function of pH. We show that Anap is highly insensitive to pH in the range 4 to 8 and it does not change fluorescence in conditions in which high outward fluxes can change the local intracellular pH. Our experiments confirm that Anap can be used without interference from local changes in proton concentration.

When incorporated at position 197, Anap produced fluorescence signals that indicate an increase in intensity with depolarization and saturated in magnitude at positive potentials. This behavior indicates dequenching of Anap as the S4 segments undergoes an outward movement during the activation conformational change. Anap has been incorporated in other membrane proteins, including the *Shaker* potassium channel, in which Anap was incorporated in the S4-S5 linker and displayed fluorescence quenching upon depolarization (Kalstrup and Blunck, 2018). Anap has been incorporated in the S4 of the hyperpolarization-activated cyclic nucleotide-gated (HCN) channel (Dai et al., 2019), where it is quenched or dequenched upon hyperpolarization in a position-dependent manner. It has also recently been incorporated at the bottom of the S4 in the voltage-dependent phosphatase, CiVSP (Mizutani et al., 2022), where it becomes quenched upon depolarization. The direction of the fluorescence changes due to S4 motion are difficult to predict, since, as we have shown, Anap’s fluorescence can be affected by both the local environment’s polarity and interaction with specific quenching groups that are part of the channel sequence.

The fluorescence changes we observe in 197Anap channels indicate that the G-V and F-V relationships have almost the same voltage-dependence at the ΔpH values tested, suggesting that S4 movement closely follows channel opening, and that S4 movement and activation of the proton conductance are equally affected by the proton gradient. A similar conclusion has been reached in studies measuring S4 movements of hH_V_1 by fluorescence (Schladt and Berger, 2020) or in *Ciona* H_V_1 by gating current recordings (Carmona et al., 2021). Interestingly, these changes in fluorescence as a function of voltage, are not accompanied by changes in the emission spectrum of Anap, suggesting that the probe remains in a solvated environment regardless of the state of the channel. This is in accordance with the finding that the VSD that forms hH_V_1 channels has a large extracellular cavity able to contain many water molecules (Ramsey et al., 2010).

Since Anap remains solvated in the closed and open state, what is the origin of the reported fluorescence changes? As is common with other fluorescent probes, we hypothesized that an aromatic residue could act as an Anap quencher and thus found that an aromatic outside the S4 close enough to have this function is F150. Mutation F150A in the 197Anap background produced an increased Anap/mCherry fluorescence intensity as compared to 197Anap alone, indicating reduced quenching. This result suggests that 197Anap moves away from F150 as the S4 segment moves outward during channel activation. It should be noted that F150A-197Anap channels still produce fluorescence changes upon depolarization. This suggests that other amino acid residues apart from F150 (aromatic, charged) can also quench Anap or changes in Anap’s quantum yield are still being produced by voltage-dependent solvent accessibility.

F150 has been shown to be part of a “hydrophobic gasket” through which S4 charges slide during channel activation. Mutations at this position and at W207 in the S4 produce altered gating (Banh et al., 2019; Cherny et al., 2015; Wu et al., 2022). In our case, F150A-197Anap shows a reduced shift of the G-V between ΔpH 0 and 1, from the expected ~40 mV to 22 mV, although the F-V curve shifts by 58 mV. These changes could be explained by altered movement of S4 through F150A, leading to changes in the coupling between voltage sensor movement and proton conductance activation. Also, this is the first time that F150 is reported as a possible actor in the pH-sensing mechanism of H_V_1.

Substitution of 1201 for Anap allowed us to uncover a slow step in the deactivation pathway. The fluorescence signal observed upon channel closure by repolarization at ΔpH = 0 shows two components, one of which is much slower than channel closing as reported by the tail current. The fact that tail current is faster than the slow component of the deactivation fluorescence signal indicates that the latter is produced by a slow intermediate transition. This experimental observation is recapitulated by a simple kinetic model. Interestingly, recordings of gating currents in mutant *Ciona* H_V_1 channels show that the charge return after depolarization can be very slow, producing gating charge immobilization (Carmona et al., 2018). This observation of a singular slow transition in hH_V_1 activation illustrates the value of fluorescence recording with a small probe such as Anap.

Our data thus far indicates that a fraction of S4 movement, as reported by the F-V relation, occurs before the increase of the proton conductance, and that S4 movement can continue after channel activation. Comparison of the V0.5 values of Q-V and G-V curves in hH_V_1 channels (De La Rosa and Ramsey, 2018) indicates that charge moves at slightly more negative values than conductance, but not at all ΔpH values. Fluorescence changes depend on all the conformational states in which the fluorophore has distinct fluorescence values, while gating currents are produced during transitions between conformations with state-dependent charge distributions (Cha and Bezanilla, 1997). For these reasons, F-V and Q-V curves of multistate channels are not expected to be identical or contain the same information.

Our fluorescence data are consistent with recent experiments that have shown that the characteristic gating effect of the proton gradient on voltage-gated proton channels comes about by a conformational change that affects voltage sensor movement and not a channel opening transition happening after voltage-sensor activation. Furthermore, our modeling suggests that all transitions in the activation pathway, including a characteristic slow transition detected by fluorescence are modulated by ΔpH.

The mechanism of ΔpH modulation is still unknown. It has been proposed that the energy stored in the pH gradient is directly coupled to S4 movement to produce ΔpH-dependent gating (Carmona et al., 2021). We have previously proposed an allosteric model in which both extracellular and intracellular protons can affect local electrostatic networks and bring about ΔpH-gating (Rangel-Yescas et al., 2021). This class of models predicts the existence of multiple open states, which is supported by the observation that S4 movement can happen after channel opening.

A mechanism in which the proton gradient energy is coupled to S4 movement predicts that the absolute value of pH should not influence gating. Interestingly, we have observed that the absolute pH values used to set up a ΔpH = 0 do affect gating. When pH_o_=5.5/pH_i_=5.5, G-V and F-V are almost superimposed and the F_off_ signal has a fast and slow component; in contrast, when pH_o_=7/pH_i_=7, the F-V curve has almost the same voltage-dependence, but conductance can be observed at more negative voltages and the F_off_ signal only contains the fast component. These results suggest that the absolute pH in the extracellular side of the channel is a determinant of the steady-state gating, presumably modulating the slow rate constant in the activation pathway.

## Materials and Methods

### Molecular biology and HEK cell expression

A plasmid containing the human voltage-gated proton channel (hH_V_1) was a gift from Dr. Ian Scott Ramsey (Virginia Commonwealth University, Richmond, VA). We used the fluorescent protein mCherry as a reporter to verify L-Anap incorporation. The construct hH_V_1-mCherry was made by the PCR overlap technique, adding the sequence of fluorescent protein mCherry after the C-terminus of hH_V_1 with the following linker sequence: (Gly-Gly-Ser)_3_. This construct was subcloned into the pCDNA3.1 vector. For all hH_V_1-TAG mutants, an amber codon (TAG) was introduced using appropriate mutagenic oligonucleotides and a protocol for whole plasmid site-directed mutagenesis employing KOD polymerase (Merck Millipore, Germany) as detailed in manufacturer’s instructions and previous work (Zheng et al., 2004; Munteanu et al., 2012). The bacterial methylated DNA templates were digested with the DpnI restriction enzyme, and the mutant plasmids were confirmed by automatic sequencing at the Instituto de Fisiología Celular, UNAM.

We used HEK293 cells for channel expression and L-Anap incorporation experiments. The HEK cells used in this study were found free of mycoplasma infection (Sigma-Aldrich mycoplasma detection kit). These cells were cotransfected with 0.1 - 1 μg of mutant hH_V_1-TAG plasmid and 0.7 μg of pAnap plasmid (a gift from Dr. Sharona Gordon, University of Washington, Seattle, WA) using the transfection reagent JetPei (Polyplus-transfection). The pANAP plasmid contains the orthogonal pair tRNA/aminoacyl tRNA synthetase specific to L-Anap. The Methyl ester form of L-Anap; L-Anap-Me (AsisChem Inc.) was added to the medium of cells in 35 mm culture dishes from a storing stock solution of 10 mM to a final concentration of 10-20 μM. Through the text, we will refer to L-Anap as Anap for simplicity. Cells were incubated during 12-48 hours before experiments in Dulbecco’s Modified Eagle Medium (DMEM, Invitrogen) supplemented with 10% fetal bovine serum (Invitrogen, USA) and penicillin-streptomycin (100 units/ml — 100 μg/ml, Invitrogen, USA) at 37 °C in a 5% CO_2_ atmosphere. Around 4 hours before electrophysiological recordings, HEK293 cells were treated with 0.05% trypsin-ethylenediaminetetraacetic acid (Trypsin-EDTA) to obtain rounded cells, which were then re-platted in 35 mm glass-bottom dishes (World Precision Instruments, USA) and used for experiments within 3-6 hrs. All the experiments were performed at room temperature (~25°C).

### Electrophysiology

Recordings of proton currents were performed in the whole-cell patch-clamp configuration using fire-polished borosilicate micropipettes (Sutter Instruments, USA). Currents were recorded by an Axoclamp 200B amplifier (Axon Instruments, USA) and acquired with an ITC-18 AD/DA converter (HEKA Elektronik, Germany), both controlled with Patchmaster software (HEKA Elektronik, Germany). Currents were low-passed filtered at 5 kHz and sampled at 20 kHz. The extracellular solution contained (in mM): 100 tetramethylammonium hydroxide and methanesulfonic acid (TMAOH-HMESO_3_), 100 buffer ((2-(N-morpholino)ethanesulfonicacid (MES) for pH 5.5, and 6.0; 4-(2-hydroxyethyl)-1-piperazineethanesulfonic acid (HEPES) for pH 7.0 and 7.5), 2 CaCl_2_, 2 MgCl_2_, 8 HCl and pH-adjusted with TMAOH and HMESO. The intracellular solution contained (in mM): 80 (TMAOH-HMESO_3_) 100 buffer (MES for pH 5.5 and 6.0; HEPES for pH 7.0 and 7.5), 10 ethylene glycol-bis(β-aminoethyl ether)-N,N,N’,N’-tetraacetic acid (EGTA), 10 MgCl_2_, and 4 HCl and pH-adjusted with TMAOH and HMESO. With these solutions, patch pipettes had a resistance of 2-5 MΩ. Since cells used in these experiments were round to improve space-clamp and currents were relatively small, no series-resistance compensation was employed. The voltage-clamp protocols varied depending on the value of ΔpH and are indicated in the figure legends. The interval between each test pulse was 45 s at the holding potential to facilitate return of slow fluorescence signals and minimize the effects of proton depletion.

### Fluorescence measurements

Fluorescence measurements in whole-cell patch-clamp fluorometry (PCF) experiments were made using a TE-2000U (Nikon, Japan) inverted epifluorescence microscope with a 60x oil immersion objective (numerical aperture 1.4). A 405 nm solid-state laser (Compass 405-50 CW, COHERENT, USA) and a filter cube containing a 405/20-nm excitation filter, a 405-nm long pass dichroic mirror, and a 425-nm long-pass emission filter were used for Anap fluorescence excitation. For mCherry fluorescence, measurements were performed using an Ar-Ion laser (Spectra-Physics, Germany) and a filter cube with a 514/10-nm excitation filter, a 514-nm long pass dichroic mirror, and a 530-nm long-pass emission filter (Chroma, USA). Both lasers were through-air coupled, collimated using an optical cage system and appropriate optics (Thorlabs, USA) and then focused into the objective’s back focal plane through the microscope’s rear port. Imaging was performed using Luca or Ixon Ultra EMCCD cameras (Andor, Oxford instruments, Ireland) controlled by Micromanager software (Edelstein et al., 2014). The fluorescence of a region without cells was measured with the same ROI employed with cells and this background was subtracted from Anap fluorescence images. Image stacks from cells were recorded at 10-25 Hz (100 - 40 ms of exposure, respectively). To improve signal-to-noise ratio, 4×4 or 8×8 pixel binning was used. Initially, fluorescence time course was measured from a region of interest (ROI) that included only the membrane of the patched cell. Identical results were obtained by using a ROI encompassing all the cell.

Fluorescence and proton current recording synchronization was achieved through a home-programed Arduino Uno microcontroller board (Arduino, Italy) triggered by a PatchMaster-generated TTL pulse.

For spectral measurements, the light from the microscope was collected by a SpectraPro 2150i imaging spectrograph (Princeton Instruments, USA) mounted between the microscope and EMCCD camera. The mCherry fluorescence was used as an indicator of membrane-associated channels. A small area from the membrane-associated mCherry fluorescence is selected using the spectrograph slit, and mCherry and Anap spectra were recorded by measuring a line scan 10 pixels wide from the cell membrane spectral image (Figure 2-Supplement 1). Background fluorescence spectrum was recorded from a region of the image without cells and subtracted from Anap and mCherry fluorescence. Unless indicated, all the spectra were obtained at resting potential in non-patched cells (Thomas and Smart, 2005a) and a ΔpH ~ −0.2.

### Data analysis

IgorPro (Wavemetrics) and ImageJ (NIH) software were used to analyze the data. For the G-V relationships, conductance (*G*) was calculated from proton currents according to:

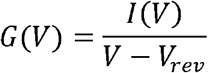

Where *V_rev_* is the proton current reversal potential, measured from the current-voltage relation. Then, *G* was normalized and fit to a Boltzmann function as follows:

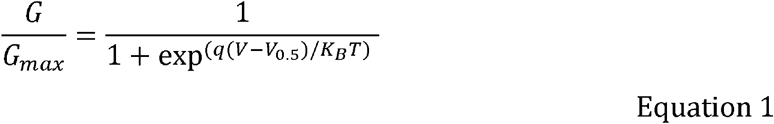

Where *q* is the apparent gating charge (in elementary charges, *e_0_*), *V* is the membrane potential, *V_0.5_* is the potential at which half of the maximal activation is reached, *K_B_* is the Boltzmann constant and *T* is the temperature in Kelvin.

The time course of fluorescence in PCF experiments, was obtained from all the background-subtracted images in a stack (*F_i_*), and the changes through time were normalized to the first image (*F_0_*) as follows:

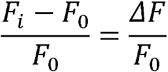

Then, this normalization was multiplied by 100 to obtain the percent change of fluorescence. This procedure was carried out for each stack at each voltage. The voltage-dependence of the fluorescence was estimated from F-V relationships. The value of the fluorescence at the end of the volage step was normalized and fit to a Boltzmann function:

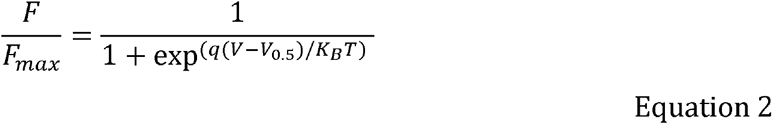

Where *F* is the percent of fluorescence change at *V* potential and *F_max_* is the maximum fluorescence percent change in each experiment at *V* potential. The meaning of *q*, *V*, *V_0.5_* and *K_B_T* is the same as in equation 1. All data are presented as the mean ± standard error of the mean (s.e.m.).

The time constants activation of proton currents and fluorescence signals were obtained by fitting of the second half of each trace to the equation:

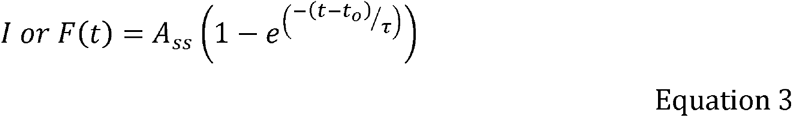

Where *A_ss_* is the amplitude of the fluorescent signal (*F*) or current (*I*) at steady state, *τ* is the time constant, and *t_o_* is the time of start if the voltage pulse, both in ms. The voltage dependence of τ was calculated from a fit to equation:

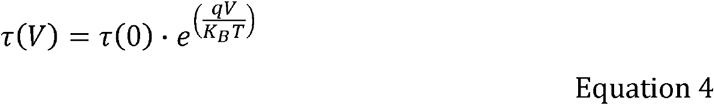

Where *τ(0)* is the activation time constant at 0 mV, *q* is the partial charge in e_0_ units and *V*, *K_B_* and *T* have the same meaning as in equation 1.

### Modelling

Modelling of current and fluorescence was carried out using custom-written programs in IgorPro (Wavemetrics). The occupancy of each discreet state in the models was calculating by numerically solving the differential equations describing the transitions between states. The occupancy of each discreet state *i* is *P_i_* and it was calculated by numerically solving the differential equations described by a master equation:

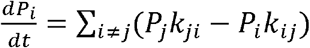

The rate constants *k_ij_* or *k_ij_* are given by:

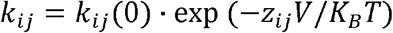

Where *k_ij_(0)* is the value of the rate constant at 0 mV, *z_ij_* the partial charge associated with the transition and *K_B_T* have the same meaning as in Eq.1.

The current as a function of time tand voltage *V* was calculated as:

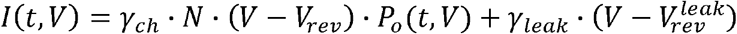

*γ_ch_* is the single proton channel conductance, *N* is the number of channels, *V_rev_* is the reversal potential and *P*_o_ is the probability of the open state. *γ_leak_* is the leak conductance and 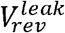 is the reversal potential of the leak currents.

The fluorescence was calculated as:

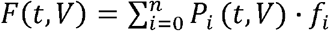

*f_i_* is the fluorescence of the *i*-th state in arbitrary units.

## Acknowledgements

We thank Eduardo Guevara for measurements of Anap spectra in different solvents and Manuel Hernández for excellent technical support. We thank Dr. Sebastian Brauchi for the loan of the 405 nm laser. This work was supported by DGAPA-PAPIIT-UNAM grant No. IN215621. E. S-D is a doctoral student from Programa de Doctorado en Ciencias Bioquímicas-UNAM and was supported by a doctoral thesis scholarship from CONACyT No. 463819 (CVU 659182). M.E.O-C. is a doctoral student from Programa de Doctorado en Ciencias Biomédicas-UNAM and is supported by a doctoral thesis scholarship from CONACyT No. 788807 (CVU 1101710)

## Notes

### Competing Interest Statement

The authors have declared no competing interest.

### Summary of Updates

New experiments have been added as well as new figures and Introduction and Discussion sections updated for clarity.

