## Supplementary data for "Activation-pathway transitions in human voltage-gated proton channels revealed by a non-canonical fluorescent amino acid"

**Supplementary figures**

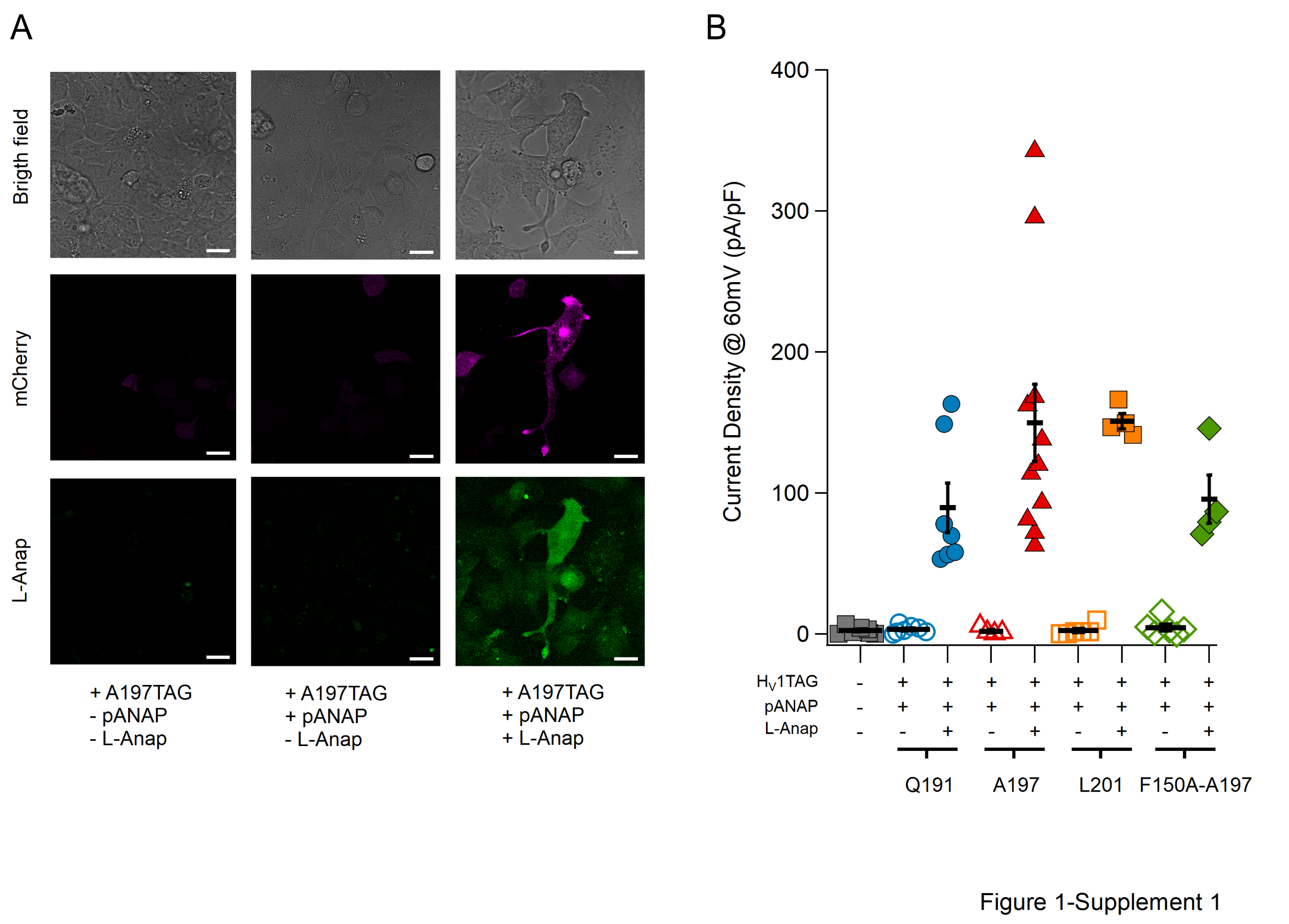

**Figure 1-Supplement 1.** L-Anap incorporation suppresses amber codons inserted in hH_V_1. A) Confocal images of HEK 293 cells in Bright field (top panel), mCherry fluorescence (middle row, magenta) and Anap fluorescence (bottom row, green). Each column shows the same field, and the bottom labels indicate transfection of the H_V_1-A197TAG plasmid in the presence or absence pAnap plasmid and L-Anap in the culture media. The images were obtained with a Leica TCS SP5 inverted confocal microscope (Leica Microsystems, Germany). Fluorescence of mCherry was excited with a He-Ne laser at 543 nm and the emission was filtered at 580-610 nm, while Anap was excited with a 405 nm laser diode, and the emission was filtered at 415-458 nm. Scale bar = 10 µm. B) Summary of current density quantification at 60 mV test pulse (ΔpH=1) in HEK293 cells co-transfected with the hH_V_1-TAG plasmid in different positions (Q191, blue circles; A197, red triangles; L201, orange squares; F150A-A197, green diamonds) and the pAnap plasmid with (+, filled markers) or without (-, empty markers) L-Anap added to incubation media. Note that the cells that weren't incubated with L-Anap presented similar current density as non-transfected cells (grey marks), showing a low probability of non-specific amino acid incorporation. Current density values: non-transfected cells: 2.6 ± 0.98 pA/pF, n=7. H_V_1-Q191TAG+pAnap: 3.4 ± 1.04 pA/pF (n=7). H_V_1-Q191TAG+pAnap+L-Anap: 89.7 ± 17.53 pA/pF (n=7). H_V_1-A197TAG+pAnap: 2.1 ± 1.2 pA/pF (n=4). H_V_1-A197TAG+pAnap+L-Anap: 149.9 ± 27.42 pA/pF (n=11). H_V_1-L201TAG+pAnap: 2.6 ± 1.47 pA/pF (n=6). H_V_1-L201TAG+pAnap+L-Anap: 151.1 ± 5.4 pA/pF (n=4). H_V_1-F150A-A197TAG+pAnap: 4.6 ± 2.4 pA/pF (n=6). H_V_1-F150A-A197TAG+pAnap+L-Anap: 95.7 ± 16.98 pA/pF (n=4). Horizontal bars and error bars represent these mean ± s.e.m. values and markers are individual experiments.

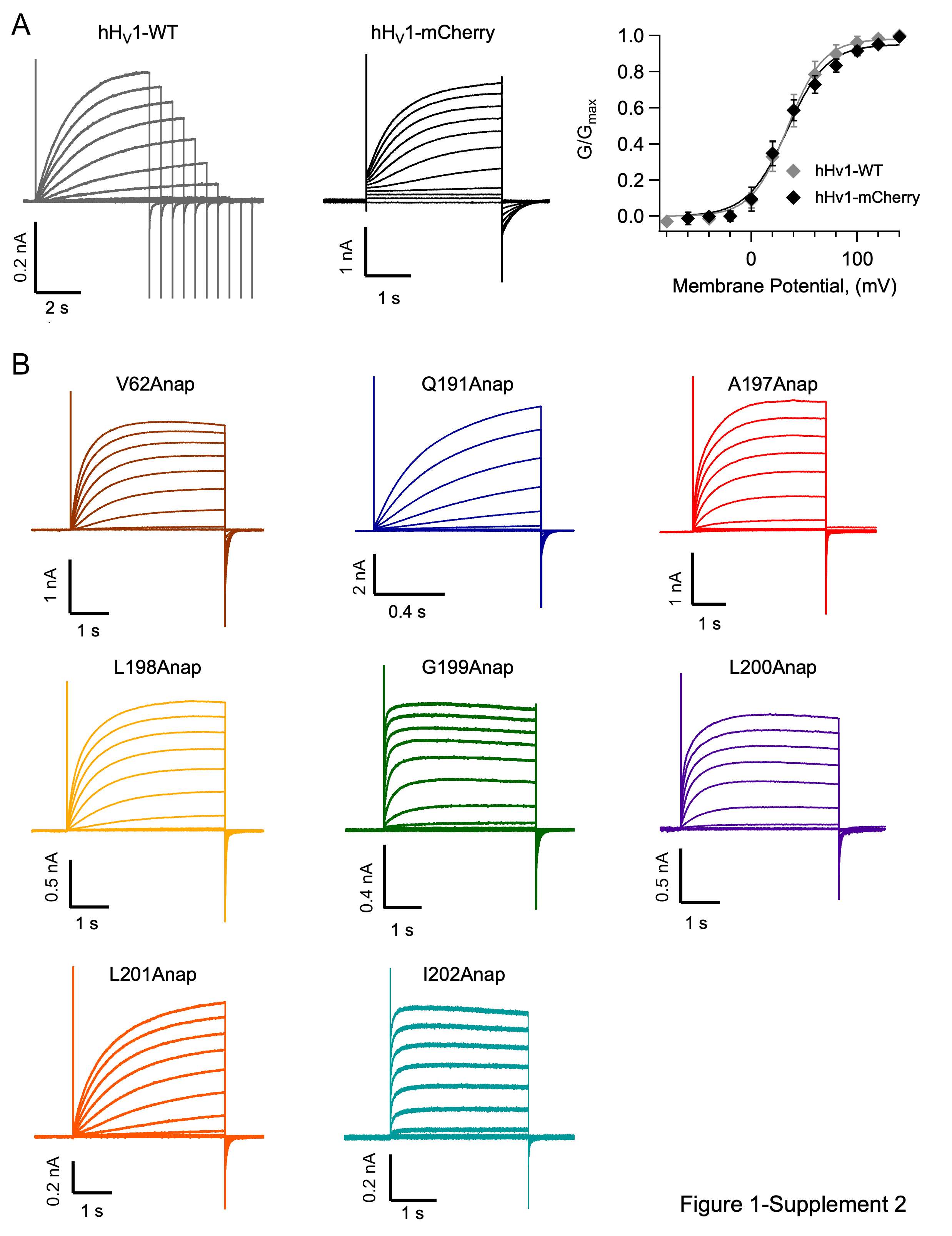
 **Figure 1-Supplement 2.** The fusion of mCherry in the C-terminus does not affect channel function and incorporating L-Anap in hH_V_1 produces functional channels. A) Family of currents of hH_V_1-WT (gray traces) and hH_V_1 with the fluorescent protein mCherry attached to channel’s C-terminus (black traces). The right panel shows the conductance vs. voltage relationship of these two constructs. Notice the similar behavior. B) Current families of channels incorporating Anap at the indicated position. Each family was obtained with a protocol of voltage pulses from –100 mV to +140 mV in 20 mV steps from a holding potential of –100 mV. Al experiments were performed at a ∆pH=1. The parameters of the Boltzmann fits are summarized in Supplementary Table I.

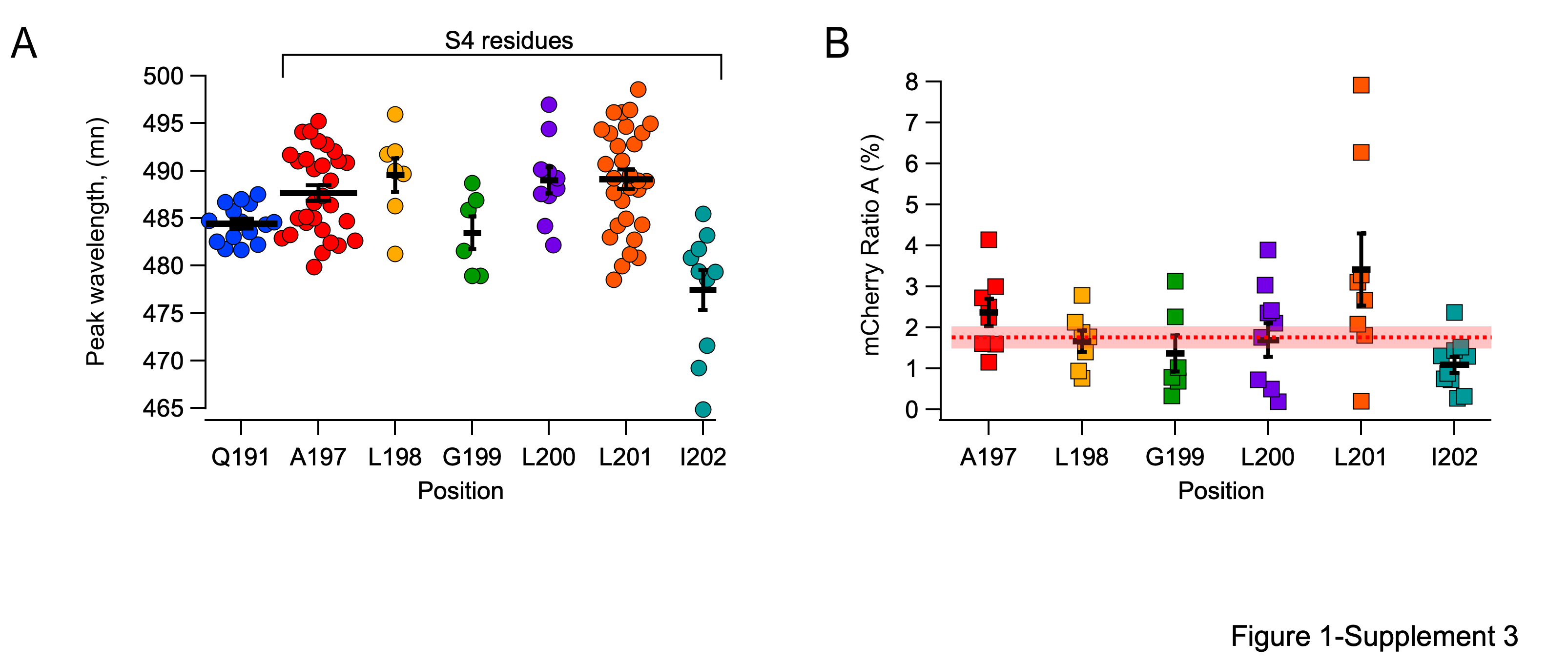

**Figure 1-Supplement 3.** Fluorescence of L-Anap attached at different hH_V_1 S4 positions. A) Measurements of L-Anap emission spectrum peak wavelength incorporated in S3-S4 loop (Q191: 484.4 ± 0.5 nm, n=15), and the sites in extracellular half of S4 studied in this work (A197: 487.7 ± 0.8 nm, n=30; L198: 489.6 ± 1.8 nm, n=7; G199: 483.5 ± 1.7 nm, n=6; L200: 489 ± 1.4 nm, n=10; L201: 489.1 ± 1 nm , n=30; I202: 477.4 ± 2.1 nm, n=10). B) Quantification of direct excitation (Ratio A) of mCherry emission spectra peak (610 nm) excited by 405 nm laser. Ratio values per position: A197: 2.4 ± 0.3 %, n=8; L198: 1.7 ± 0.3 %, n=7; G199: 1.4 ± 0.4 %, n=6; L200: 1.7 ± 0.4 %, n=10; L201: 3.4 ± 0.9 %, n=8; I202: 1.1 ± 0.3 %, n=10. The horizontal red dotted line is the ratio A of WT-hH_V_1-mCherry (1.87 %) and the shaded area is the s.e.m. In both figures the horizontal bars and error bars represent these mean ± s.e.m. values and markers are individual experiments.

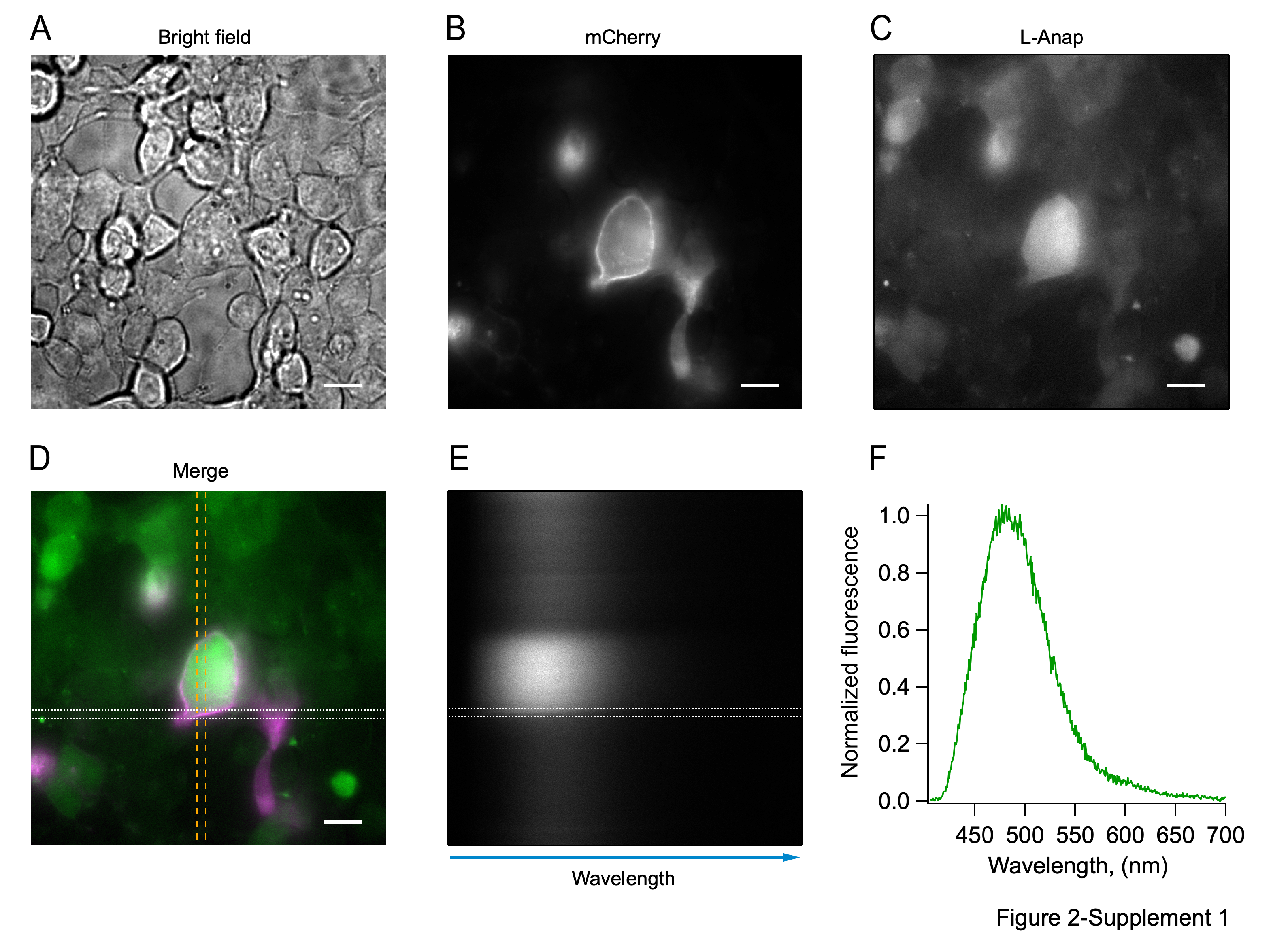

**Figure 2-Supplement 1.** Procedure for Anap spectrum measurement in cells subjected to L-Anap incorporation conditions (pAnap + H_V_1TAG + L-Anap). The HEK293 cell to be recorded is chosen in the bright field (A) and exciting the fluorescent signal of mCherry (B) at 514 nm. When the L-Anap fluorescence is confirmed, exciting the cell at 405 nm (C), a portion of the cellular membrane is isolated with the spectrograph slit (D, discontinuous vertical yellow lines; mCherry signal in magenta; L-Anap signal in green). The spectrograph scatters the L-Anap light that goes through this slit (E), producing a spectral image and a line scan is performed (white discontinuous horizontal lines in E and D) from the membrane identified by mCherry fluorescence. The fluorescence intensity measured in this scan is presented in F. Scale bar = 10 µM.

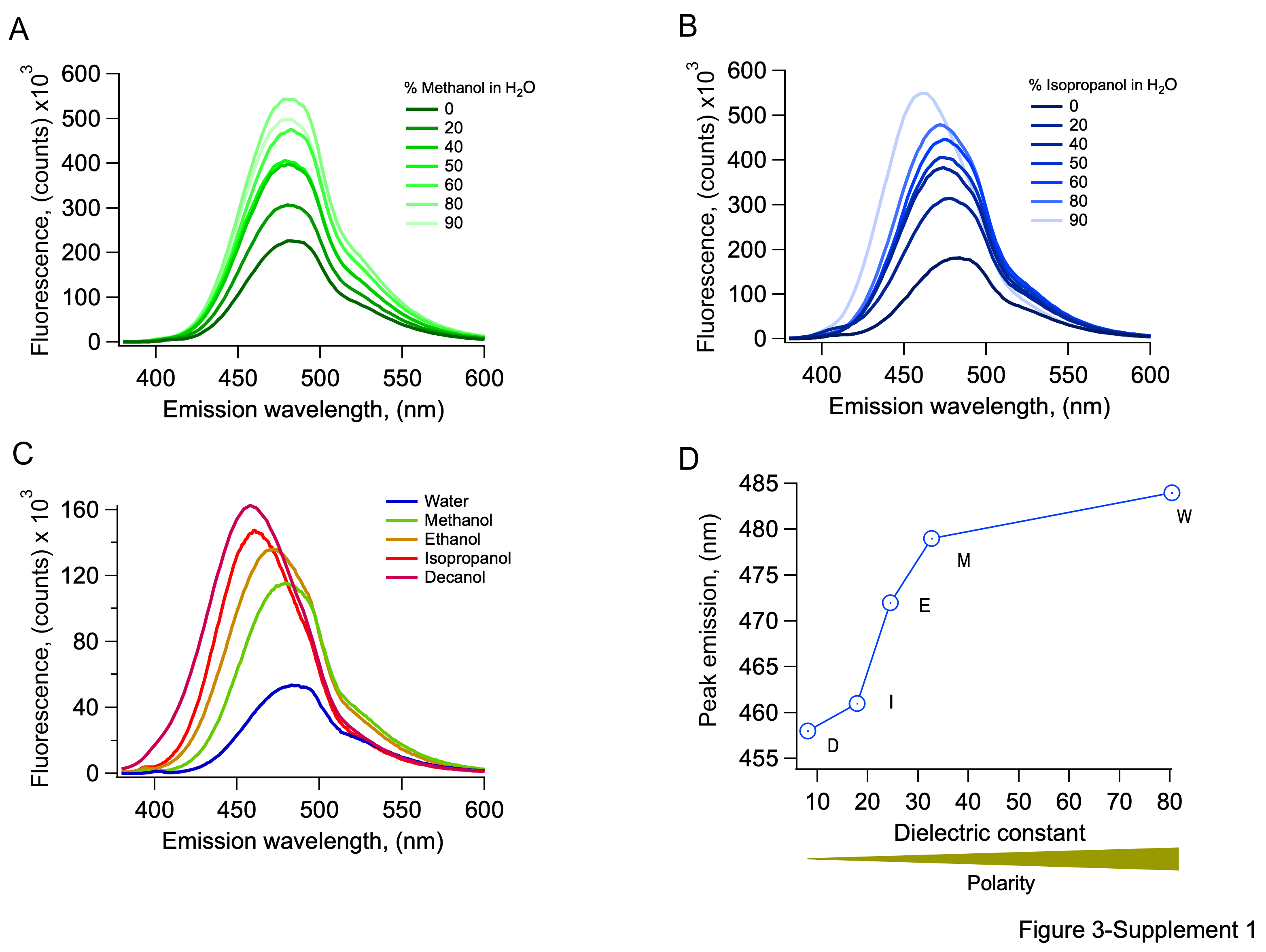
 **Figure 3-Supplement 1.** Properties of L-Anap fluorescence in solutions of solvents of different polarities. A) Emission spectrum of L-Anap in methanol-water mixtures. The percentage of methanol is indicated in the figure. Increasing methanol concentrations increase the fluorescence intensity and produce a blue shift in the peak emission wavelength. B) Emission spectrum of L-Anap in isopropanol-water mixtures. As with methanol, the percentage of isopropanol is indicated in the figure. Increasing concentrations of isopropanol also increase fluorescence intensity and a more evident blue shift in the peak emission wavelength. C) Comparison of emission spectra of L-Anap in pure solvents. Lower polarity solvents produce an increased fluorescence intensity. D) The peak emission wavelength of L-Anap obtained from spectra as in C is red-shifted at higher dielectric constants (higher polarity). All spectra were measured in a UV-VIS spectrofluorometer (PC1, ISS, USA). The excitation wavelength was 360 nm.

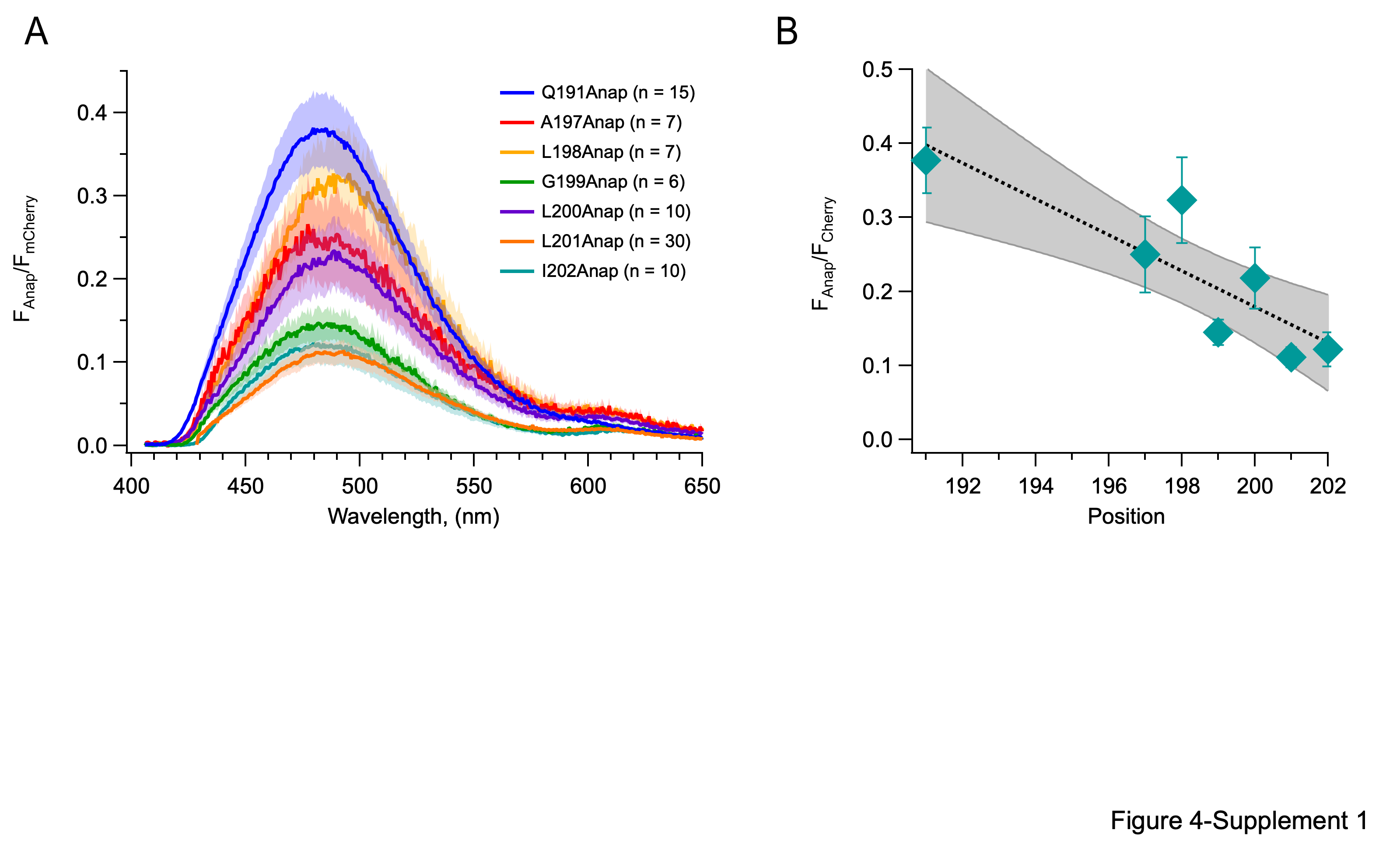

**Figure 4-Supplement 1.** The fluorescence of Anap is diminished as it is incorporated in residues closer to the center of the S4 segment. A) Fraction of Anap fluorescence in relation to mCherry fluorescence determined from emission spectra. Data are averaged spectra; the shaded regions are ± s.e.m. B) The Anap/mCherry fluorescence ratio measured at the peak emission wavelength for each construct as a function of incorporation site position. The dotted line is a linear fit and the shaded area is the 96 % confidence interval. Data are mean ± s.e.m.

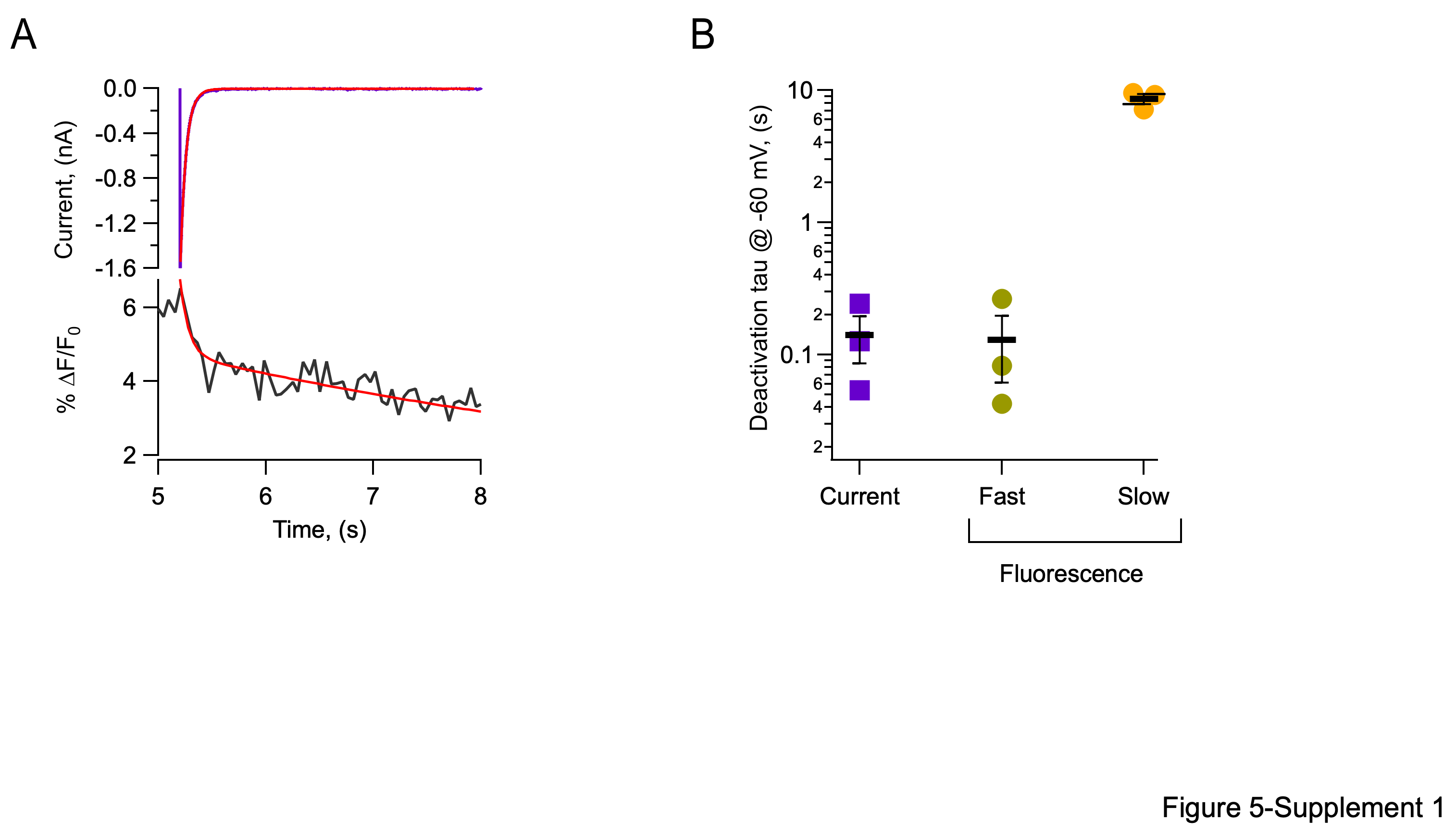

**Figure 5-Supplement 1.** Kinetics of the OFF-fluorescence signals from L201Anap channels. A) Tail current (purple trace) with a superimposed exponential fit (red). Simultaneous fluorescence decay signal (black) with a double exponential fit (red). Both recordings were obtained at -60 mV after a test pulse to 160 mV and a ∆pH =0 (pH_i_=5.5/pH_o_=5.5). B) Summary of time constants obtained from traces as in (A). The horizontal bars and error bars represent mean ± s.e.m. values and markers are individual experiments.

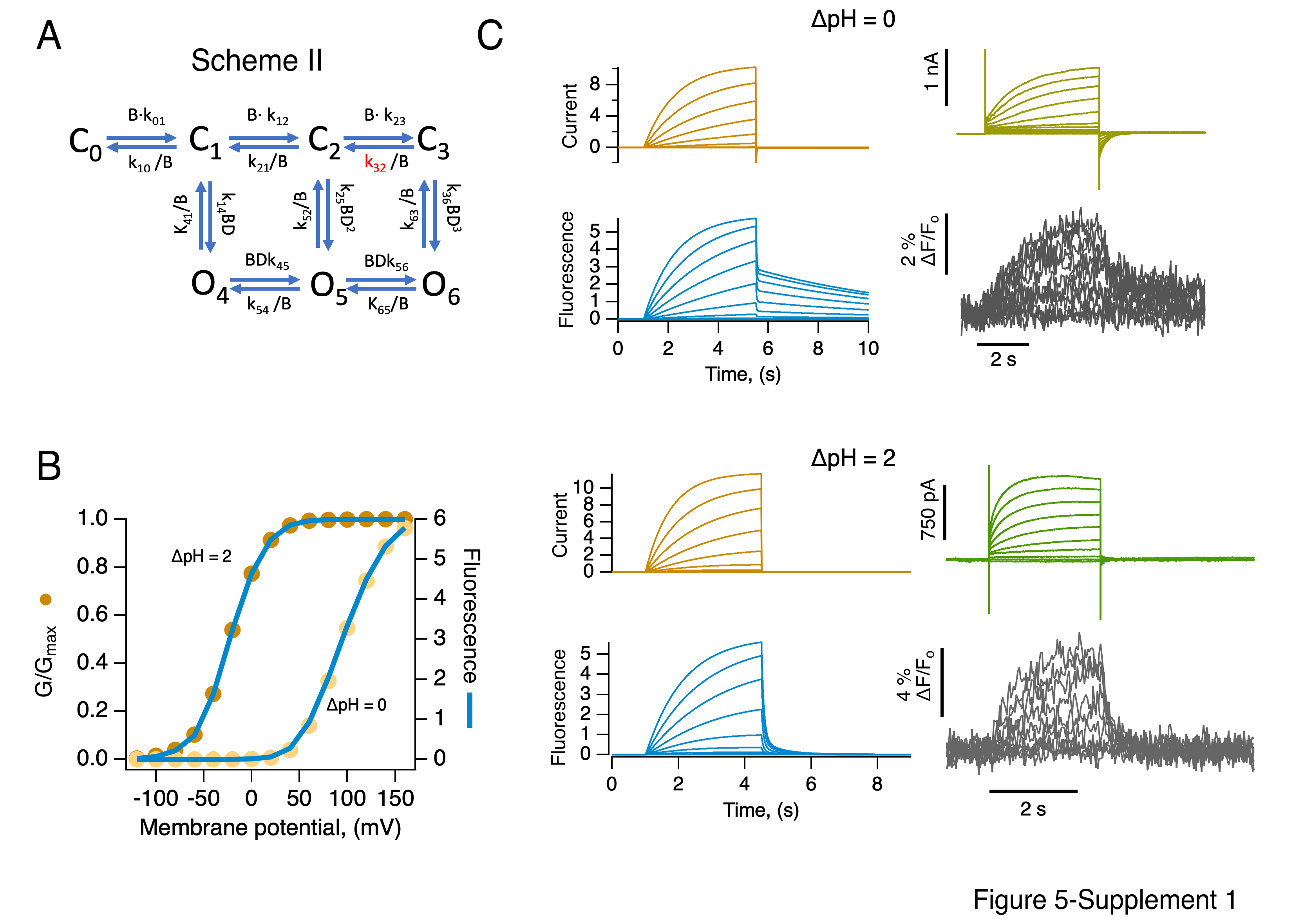
 **Figure 5-Supplement 2.** An allosteric model can explain the kinetics and voltage-dependence of fluorescence and conductance. A) Allosteric model of H_V_1 activation by voltage. The k_ij_‘s are rate constants and D is an allosteric coupling factor for channel opening. B is an empirical factor that accounts for the pH-dependence of the rate constants. In this model, channel opening can occur without full voltage-sensor movement. The closing rate constant indicated in red has a value much smaller than all other (at 0 mV) and is responsible for the slow return of the fluorescence signal upon repolarization. B) Simulation comparing the steady-state conductance and fluorescence as a function of voltage. In this mechanism, fluorescence appears at the same voltages as conductance since channel opening can occur from early closed states and currents can flow without full voltage sensor activation. C) Comparison of simulated currents (orange traces) and fluorescence signals (blue traces) with experimentally recorded currents (green) and fluorescence (grey) at the two indicated values of ∆pH, using the parameters given in Supplementary Table III.

**Supplementary Table I**. Boltzmann equation fit parameters of each mutant at ∆pH=1

| Mutant | V_0.5_ (mV) | e_0_ |
| --- | --- | --- |
| hH_V_1-WT | 34.1 ± 1.5 | 1.5 ± 0.1 |
| hH_V_1-mCherry | 33.8 ± 2.5 | 1.3 ± 0.1 |
| V62Anap | 24.4 ± 1.6 | 1.6 ± 0.1 |
| Q191Anap | 40.9 ± 4.4 | 1.4 ± 0.3 |
| A197Anap | 23.4 ± 1.1 | 1.5 ± 0.1 |
| L198Anap | 15.2 ± 1.1 | 1.8 ± 0.1 |
| G199Anap | 31.3 ± 0.7 | 1.5 ± 0.1 |
| L200Anap | 10.2 ± 2.2 | 2.4 ± 0.4 |
| L201Anap | 6.3 ± 2.2 | 1.2 ± 0.1 |
| I202Anap | -31.8 ± 0.8 | 1.7 ± 0.1 |
| F150A-A197Anap | 22.7 ± 2.3 | 0.9 ± 0.1 |

**Supplementary table II.** Parameters used in the fits to the data in Figure 5d of the model in Scheme I

|  | ∆pH =0 | | ∆pH =2 | |  |  |
| --- | --- | --- | --- | --- | --- | --- |
| Rate constant | @ 0 mV, (s^-1^) | Partial charge, (e_o_) | @ 0 mV, (s^-1^) | Partial charge, (e_o_) | State | Fluorescence, (a.u.) |
| k01 | 0.15 | 0.3 | 0.8 | 0.3 | 0 | 0 |
| k10 | 5 | -0.3 | 5 | -0.3 | 1 | 1 |
| k12 | 2 | 0.4 | 16 | 0.4 | 2 | 2 |
| k21 | **0.06** | -0.3 | **1** | -0.3 | 3 | 4 |
| k23 | 0.1 | 0.8 | 0.88 | 0.8 |  |  |
| k32 | 0.5 | -0.8 | 1 | -0.8 |  |  |

**Supplementary Table III.** Parameters used in the simulations with Scheme II shown in Figure 5 Supplement 2.

|  | ∆pH =0 | | ∆pH =2 | |  |  |
| --- | --- | --- | --- | --- | --- | --- |
| Rate constant | @ 0 mV, (s^-1^) | Partial charge, (e_o_) | @ 0 mV, (s^-1^) | Partial charge, (e_o_) | State | Fluorescence, (a.u.) |
| k01 | 0.1*B | 0.5 | 0.1*B | 0.4 | 0 | 0 |
| k10 | 1/B | -0.3 | 1/B | -0.3 | 1 | 0 |
| k12 | 0.5*B | 1.1 | 0.5*B | 1.1 | 2 | 0 |
| k21 | 0.5/B | -0.4 | 0.5/B | -0.4 | 3 | 3 |
| k23 | 2*B | 0.6 | 2*B | 0.6 | 4 | 5 |
| k32 | **0.02/B** | -0.3 | **200/B** | -0.3 | 5 | 6 |
| k14 | 1*B*D | 0.5 | 1*B*D | 0.5 | 6 | 6 |
| k41 | 3/B | -0.6 | 3/B | -0.6 |  |  |
| k25 | 1*B*D^2^ | 0.5 | 1*B*D^2^ | 0.5 |  |  |
| k52 | 3/B | -0.6 | 3/B | -0.6 |  |  |
| k36 | 1*B*C | 0.5 | 1*B*C | 0.5 |  |  |
| k63 | 3/B | -0.6 | 3/B | -0.6 |  |  |
| k45 | 1*B*D^3^ | 1.1 | 1*B*D^3^ | 1.1 |  |  |
| k54 | 3/B | -0.4 | 3/B | -0.4 |  |  |
| k56 | 1*B | 0.6 | 1*B | 0.6 |  |  |
| k65 | 0.1/B | -0.3 | 0.1/B | -0.3 |  |  |
| B | **0.33** |  | **5** |  |  |  |
| D | 2 |  | 2 |  |  |  |

Parameter with values labeled in bold indicate that these are ∆pH-dependent.
